# Cdc73 protects Notch-induced T-cell leukemia cells from DNA damage and mitochondrial stress

**DOI:** 10.1101/2023.01.22.525059

**Authors:** Ashley F. Melnick, Anna C. McCarter, Shannon Liang, Yiran Liu, Qing Wang, Nicole A. Dean, Elizabeth Choe, Nicholas Kunnath, Geethika Bodanapu, Carea Mullin, Fatema Akter, Karena Lin, Brian Magnuson, Surinder Kumar, David B. Lombard, Andrew G. Muntean, Mats Ljungman, JoAnn Sekiguchi, Russell J.H. Ryan, Mark Y. Chiang

**Affiliations:** Cellular and Molecular Biology Program, University of Michigan School of Medicine, Ann Arbor, MI; Department of Anesthesiology, Perioperative and Pain Medicine, Stanford University, Stanford, CA; Division of Hematology-Oncology, Department of Internal Medicine, University of Michigan School of Medicine, Ann Arbor, MI; Cancer Biology Program, Stanford University, Stanford, CA; Cancer Biology Program, University of Michigan School of Medicine, Ann Arbor, MI; Department of Computational Medicine and Bioinformatics, University of Michigan, Ann Arbor, MI; Center for Healthcare Outcomes and Policy, University of Michigan School of Medicine, Ann Arbor, MI; Boston University Chobanian & Avedisian School of Medicine, Boston, MA; Michigan Center for Translational Pathology, University of Michigan School of Medicine, Ann Arbor, MI; Department of Pathology and Laboratory Medicine and Sylvester Comprehensive Cancer Center, Miller School of Medicine, University of Miami, Miami, FL; Department of Pathology, University of Michigan, Ann Arbor, MI; Department of Radiology Oncology, University of Michigan School of Medicine, Ann Arbor, MI; Department of Human Genetics, University of Michigan School of Medicine, Ann Arbor, MI.

## Abstract

Activated Notch signaling is highly prevalent in T-cell acute lymphoblastic leukemia (T-ALL) but pan-Notch inhibitors were toxic in clinical trials. To find alternative ways to target Notch signals, we investigated Cell division cycle 73 (Cdc73), which is a Notch cofactor and component of transcriptional machinery, a potential target in T-ALL. While we confirmed previous work that CDC73 interacts with NOTCH1, we also found that the interaction in T-ALL was context-dependent and facilitated by the lymphoid transcription factor ETS1. Using mouse models, we showed that Cdc73 is important for Notch-induced T-cell development and T-ALL maintenance. Mechanistically, Cdc73, Ets1, and Notch intersect chromatin at promoters and enhancers to activate oncogenes and genes that are important for DNA repair and oxidative phosphorylation. Consistently, *Cdc73* deletion in T-ALL cells induced DNA damage and impaired mitochondrial function. Our data suggests that Cdc73 might promote a gene expression program that was eventually intersected by Notch to mitigate the genotoxic and metabolic stresses of elevated Notch signaling. We also provide mechanistic support for testing inhibitors of DNA repair, oxidative phosphorylation, and transcriptional machinery. Inhibiting pathways like Cdc73 that intersect with Notch at chromatin might constitute a strategy to weaken Notch signals without directly targeting the Notch complex.

## Introduction

Activation of the Notch signaling pathway plays essential roles in development and homeostasis of diverse tissues, including multiple stages of T-cell fate specification and development (Siebel and Lendahl 2017). In normal cells, the Notch1-4 receptors are activated by ligands. Next, gamma-secretase cleaves Notch, which releases <u>I</u>ntra<u>C</u>ellular <u>N</u>otch (ICN1-4). ICN translocates to the nucleus where it binds its DNA-binding cofactor RBPJ to induce transcription (Kopan 2012). In T-cell acute lymphoblastic leukemia (T-ALL), Notch signaling is activated constitutively to supraphysiological levels through *NOTCH1* mutations that occur in ~60% of cases (Weng et al. 2004). NOTCH1 cleavage is diffuse throughout the tumor (Kluk et al. 2013). Gamma-secretase inhibitors (GSIs) inhibit Notch activation in both normal and cancer cells. Early clinical studies reported excessive toxicities with continuous GSI dosing (Krop et al. 2012; Tolcher et al. 2012). To circumvent toxicity, we and others investigate the possibility of targeting transcriptional regulators that intersect with the Notch pathway since leukemia associated *NOTCH1* alleles are inherently weak transactivators (Chiang et al. 2008). Proteomic studies show that the core Notch complex can interact with transcriptional regulators that co-bind its response elements in T-ALL (Yatim et al. 2012; Lin et al. 2013; Pinnell et al. 2015) and in other cellular contexts (Borggrefe and Liefke 2012; Bray and Gomez-Lamarca 2018). In theory, inhibiting these “Notch cofactors” might interfere with oncogenic Notch signals without disrupting all Notch functions.

The polymerase-associated factor 1 complex (Paf1C) was previously linked to the Notch pathway as *Drosophila* with mutations in *Rtf1* (a Paf1C subunit) or *Bre1* (a direct Paf1C cofactor) show notches in their wings and impaired transcription of Notch target genes (Bray et al. 2005; Tenney et al. 2006). Paf1C acts as a scaffold that physically links mRNA transcriptional machinery (e.g. RNA polymerase II or Pol II) to chromatin modifying enzymes. It promotes deposition of histone marks associated with transcriptional activation (H2BK120ub, H3K4me, H3K79me, and H3K36me) and mRNA synthesis from initiation and elongation to termination and processing (Jaehning 2010; Van Oss et al. 2017; Francette et al. 2021). Paf1C is associated with active chromatin at gene bodies but is not a universal inducer of transcription (Chen et al. 2015). In its canonical role, Paf1C is estimated to promote mRNA synthesis of ~15-20% of the most highly expressed genes (Penheiter et al. 2005; Yu et al. 2015). In contrast, Paf1C has non-canonical functions at enhancers, regulating eRNA synthesis and H3K27ac deposition (Chen et al. 2017; Ding et al. 2021). CDC73 is a subunit of Paf1C that was previously showed to interact with NOTCH1 in breast cancer cells (Kikuchi et al. 2016). However, the precise link between CDC73 and NOTCH1 and its relevance for Notch-driven cancers remains unclear. Investigating CDC73 is also attractive since T-ALL cells are known to be “addicted” to components of transcriptional machinery (Bradner et al. 2017). In this context, we proposed and experimentally tested the role of CDC73 in Notch-induced T-ALL. Our results show that CDC73 intersects Notch-regulated pathways, particularly in DNA repair and oxidative phosphorylation, through its canonical functions and suggest CDC73 as a potential therapeutic target in this cancer.

## Results

### CDC73 interacts with NOTCH1 and ETS1 in T-ALL cells

Previous studies showed that CDC73 and associated factors interact with NOTCH1 and enhance Notch target gene expression (Bray et al. 2005; Tenney et al. 2006; Kikuchi et al. 2016). To confirm this interaction in T-ALL cells, we performed co-immunoprecipitation (co-IP) assays. We detected endogenous CDC73 with Flag-NOTCH1 pulldown but were unable to detect NOTCH1 with reciprocal pulldown (Fig. S1A). Since we previously showed that the transcription factor ETS1 nucleates multiple transcription factors at chromatin, including NOTCH1 (McCarter et al. 2020), we wondered if CDC73 had a stronger interaction with ETS1 than NOTCH1. Consistently, ETS1 interacted with CDC73 in forward and reciprocal co-IP assays with Flag-tagged proteins (Fig. S1A-B) and endogenous proteins (Fig. S1C). To define the genomic locations of CDC73-associated complexes, we performed CDC73 ChIP-Seq in control and ETS1 knockdown in the T-ALL cell line THP-6 and integrated these datasets with our previously published RNA-Seq and ChIP-Seq datasets (McCarter et al. 2020). Consistent with a previous report of PAF1 (a PAF1C member) in myeloid leukemia cells (Yu et al. 2015), the strongest CDC73 signals were associated with the most highly transcribed genes (Fig. S1D) and active chromatin (Fig. S1E). Consistent with our co-IP data, CDC73 peaks overlapped more frequently with ETS1 peaks (56%) than RBPJ/Notch peaks (40%) (Fig. S1F). The majority of RBPJ peaks that overlapped with CDC73 peaks did so in the presence of ETS1 (86%). Further, motif analysis showed that CDC73 peaks were more frequently associated with ETS motifs than any other motif (Fig. S1G). ETS1 knockdown attenuated CDC73 peaks that overlapped with dynamic ETS1 peaks (defined as ETS1 peaks that were significantly diminished by ETS1 knockdown; Fig. S1H-I) relative to all CDC73 peaks. Thus, while we confirmed previous work from others that NOTCH1 does interact with CDC73, the interaction appears to be dependent, at least partially, on an ETS1 context.

### Cdc73 is important for Notch-dependent T-cell development

Paf1C is important for Notch-dependent wing development in *Drosophila* (Bray et al. 2005; Tenney et al. 2006). To investigate the importance of Cdc73 for Notch functions in T cells, we first studied its effect in T-cell development, which is driven primarily by Notch signals (Hosokawa and Rothenberg 2021). Murine Notch-dependent T-cell development proceeds in the thymus through a series of stages from the double-negative (DN) stages (DN1-DN4) to the immature single positive (ISP) and CD4^+^CD8^+^ double-positive (DP) stages, and then to the single-positive (SP) CD4^+^ or CD8^+^ stages. Notch1 and Ets1 are essential for the transition between the DN stage to the DP stage (Wolfer et al. 2002; Eyquem et al. 2004; Tanigaki et al. 2004; Maillard et al. 2006). Like Notch1 and Ets1, Cdc73 and other Paf1C members are expressed throughout T-cell development (Fig. S2A-B). To test whether Cdc73 might have similar functions as Notch1 and Ets1 in early T cells, we crossed *LckCre* mice with *Cdc73^f/f^* mice (Wang et al. 2008) to generate *LckCre Cdc73^f/f^* mice (*Cdc73^Δ/Δ^* mice). Like Notch-deficient and Ets1-deficient mice, *Cdc73^Δ/Δ^* mice showed severely impaired thymopoiesis (Fig. 1A-B) with significant loss of early T cells by the DN4 stage (Fig. 1C-H) and impaired DN-to-DP cell transition (Fig. 1I-O). Expression analysis of sorted DN subsets showed that *Cdc73* deletion was visible at the DN3a stage but was strongest at DN4 stage (Fig. 1P). This effect is consistent with DN4 showing the most significant loss of cell number (Fig. 1H) and most significant loss of expression of classic Notch target genes *Hes1* and *Myc* (Fig. 1Q-R). Thus, like *Notch1* and *Ets1*, *Cdc73* promotes the DN-to-DP transition.

**Figure 1.**
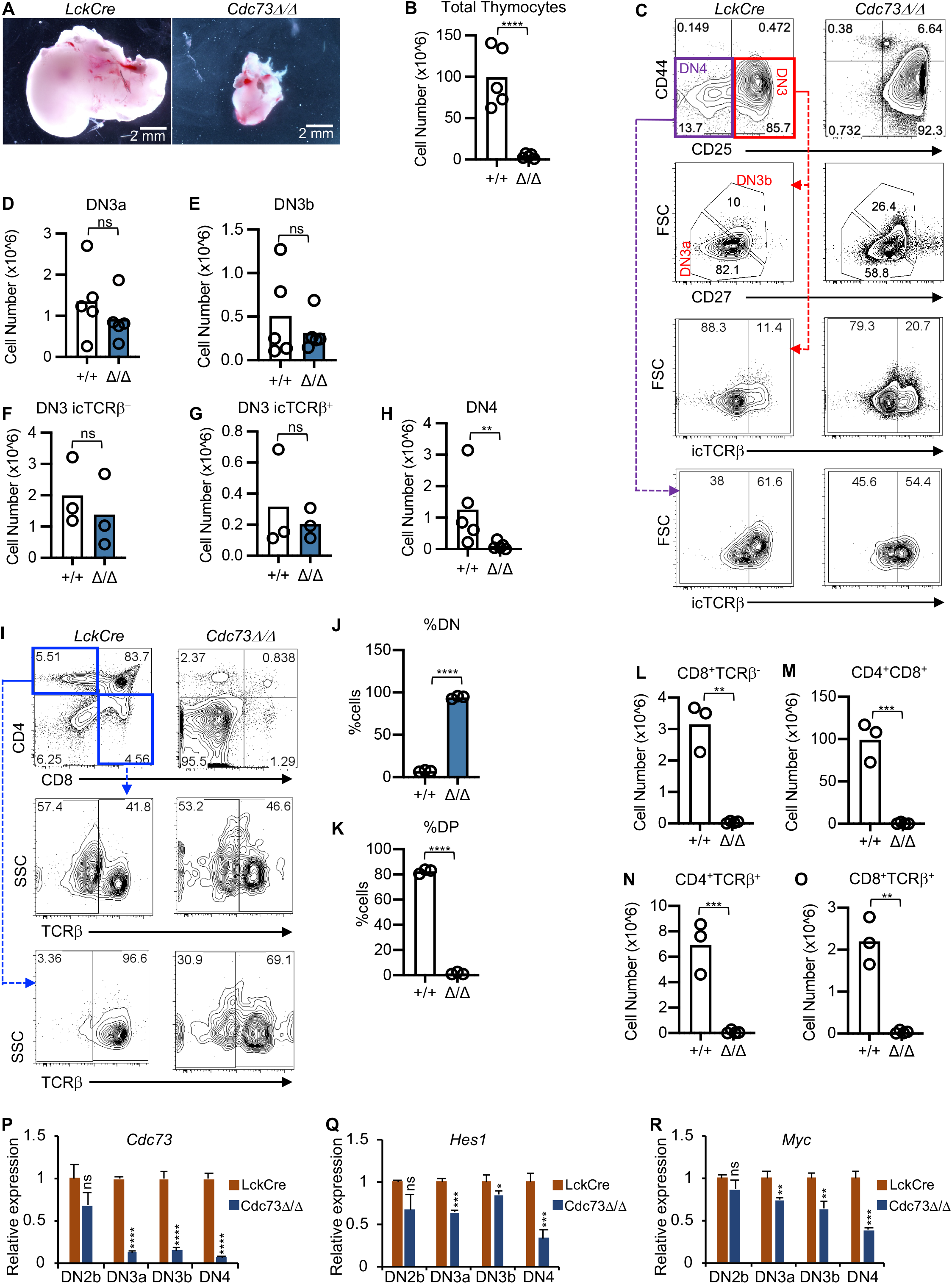
Cdc73 is important for Notch-dependent T-cell development. A-H) Representative images of thymuses (A); absolute thymocyte counts (B); representative flow cytometric profiles of DN subsets (C); and absolute numbers of DN3a (D), DN3b (E), DN3 icTCRβ^−^(F), DN3 icTCRβ^+^ (G), and DN4 (H) subsets in *LckCre* control and *LckCre Cdc73^f/f^ (Cdc73^Δ/Δ^)* mice. I-O) Representative flow cytometric profiles of CD4/CD8 subsets (I); %DN (J); %DP (K); and absolute numbers of ISP (L), DP (M), CD4 SP (N), and CD8 SP (O) thymic subsets in *LckCre* control and *Cdc73^Δ/Δ^* mice. DN3a=Lineage^−^CD44^−^CD25^+^FSC^lo^CD27^−^. DN3b=Lineage^−^CD44^−^CD25^+^FSC^hi^CD27^+^. DN4=Lineage^−^CD44^−^CD25^−^. ISP=CD8^+^TCRβ^−^. DP=CD4^+^CD8^+^. CD4 SP=CD4^+^TCRβ^+^. CD8 SP=CD8^+^TCRβ^+^. ^*^P<0.05; ^**^P<0.01; ^***^P<0.001; ^****^P<0.0001. P-R) Relative expression of *Cdc73* (P), *Hes1* (Q), and *Myc* (R) in sorted thymic subsets from *LckCre mice* (red) and *Cdc73^Δ/Δ^* mice (blue).

### Cdc73 is important for murine Notch-induced T-ALL maintenance

Next, we wondered whether the dependence of T-cell precursors on *Cdc73* would be conserved after they transform to leukemia. To test this possibility, we used a well-established murine model of Notch-induced T-ALL (Pear et al. 1996; Aster et al. 2000). We transduced bone marrow stem and progenitor cells from *Rosa26CreER^T2^* or *Rosa26CreER^T2^ Cdc73*^f/f^ mice with an activated *Notch1* allele (*ΔE/Notch1*) (Schroeter et al. 1998; Weng et al. 2003). We transplanted these cells into recipient mice to generate primary tumors (Fig. 2A, S3A). Next, we transplanted primary tumors into secondary recipients, which were injected with tamoxifen to delete *Cdc73*. In contrast to control T-ALL mice (Fig. 2B), *Cdc73^f/f^* T-ALL mice treated with tamoxifen showed reduced blast or white blood cell counts of 700-fold or 11-fold respectively (Fig. 2C). Tamoxifen did not change median survival relative to vehicle in control T-ALL mice (Fig. 2D) but prolonged survival by >200% or 84% in *Cdc73^f/f^* T-ALL mice (Fig. 2E). Thus, Cdc73 is important for maintenance of Notch-induced T-ALL.

**Figure 2.**
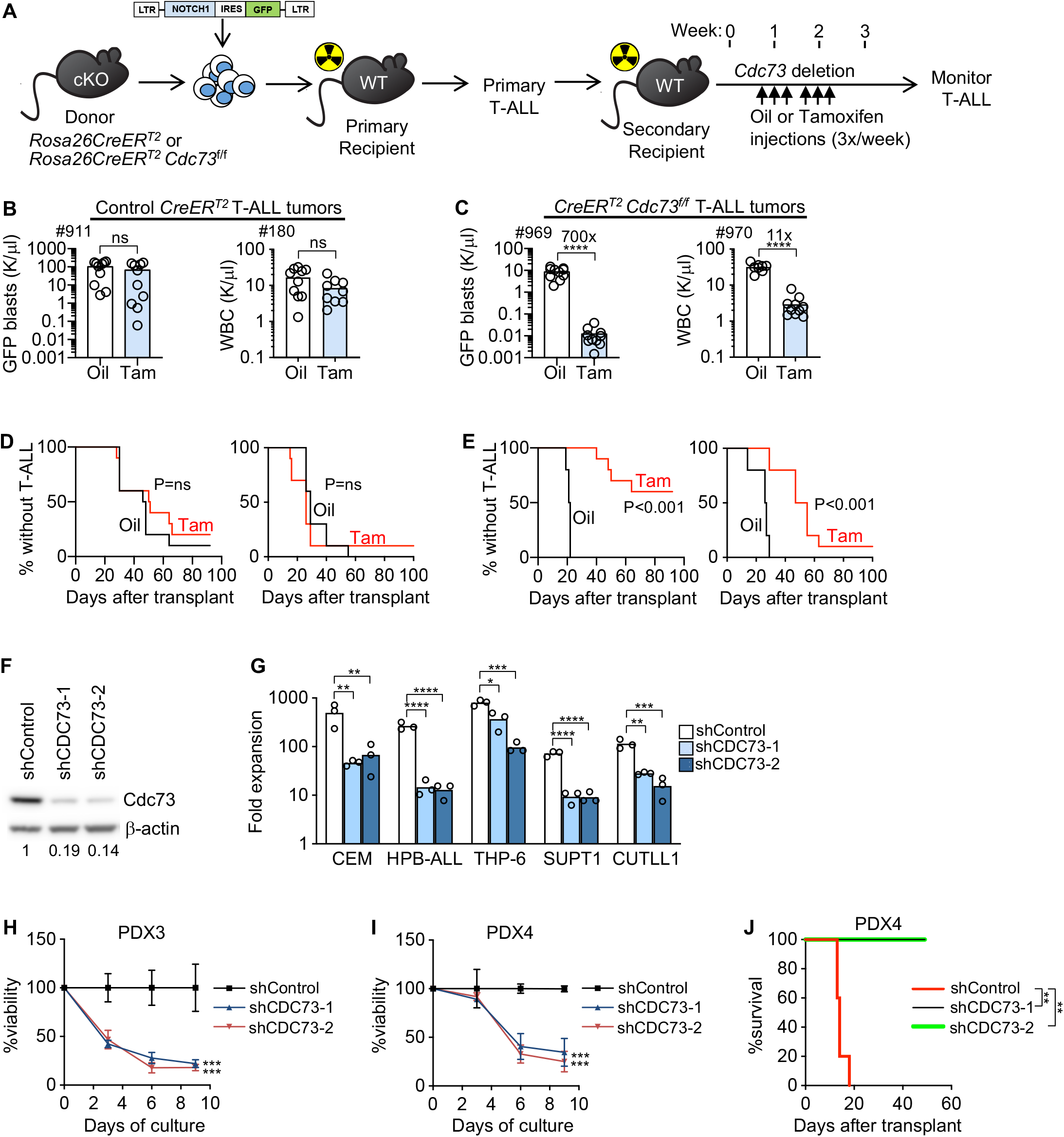
Cdc73 is important for Notch-induced T-ALL maintenance. A) Experimental strategy to study dependence of Notch-induced T-ALL maintenance on *Cdc73*. Tam=25mg/kg tamoxifen. B-E) Mice were injected with 2 independent *ΔE/Notch1*-induced *Rosa26CreER^T2^* control T-ALL tumors (B, D) or 2 independent *ΔE/Notch1*-induced *Rosa26CreER^T2^ Cdc73^f/f^* murine T-ALL tumors (C, E). Numbers indicate tumor IDs. Peripheral blood GFP^+^ counts or WBC (B-C) at 2.5 weeks post-transplant and survival (D-E) were measured. F) Western blot showing *CDC73* knockdown in shRNA-transduced CEM cells. Numbers indicate relative band intensity. G) Fold expansion (Day 9 cell count/Day 0 cell count) of Notch1-activated T-ALL cells transduced with two independent shCDC73. H-I) Viability of conventional T-ALL PDX cells transduced with shCDC73 in OP9-DL4 stromal cell culture. N=3. J) Leukemia-free survival of NSG mice injected with PDX4 cells transduced with shCDC73 that were passaged in NSG mice for 24 weeks. N=5. ^**^P<0.01; ^***^P<0.001; ^****^P<0.0001.

### CDC73 is important for human *NOTCH1*-activated T-ALL propagation and maintenance

In a clinically annotated cohort of pediatric T-ALL, *CDC73* is expressed across all oncogenomic and developmental subgroups (Fig. S3B-C). *CDC73* expression is not associated with WBC (Fig. S3D) or survival (Fig. S3E). To test the functional importance of *CDC73* for human T-ALL cell proliferation, we transduced *CDC73* shRNAs into *NOTCH1*-activated, ETS1-dependent T-ALL cells with effective suppression of CDC73 protein (Fig. 2F). *CDC73* knockdown reduced proliferation of NOTCH1-activated T-ALL cell lines by 2-9-fold (Fig. 2G). To test the anti-tumor effects of *CDC73* inactivation in non-immortalized human T-ALL cells, we took advantage of the success of shRNA protocols in knocking down gene expression in Notch-activated patient-derived xenografts (PDXs) (Yost et al. 2013; McCarter et al. 2020). *CDC73* knockdown reduced viability of PDX cells by 5 or 6-fold (Fig. 2H-I). Transplantation of these cells into immunodeficient NSG mice showed that *CDC73* knockdown significantly prolonged survival (Fig. 2J). Taken together, these results demonstrate strong and highly prevalent *CDC73*-dependency in human NOTCH1-activated T-ALL.

### Cdc73 promotes gene expression in DNA repair and oxidative phosphorylation (OXPHOS) pathways

Next, we sought to understand the underlying mechanism that explains why Cdc73 is required for Notch1-activated T-ALL maintenance. Towards this goal, we generated two independent Notch-induced *CreER^T2^ Cdc73^f/f^* T-ALL cell lines (969 and 970) by transferring the tumors described in Fig. 2C to cell culture media. We confirmed that 4-hydroxytamoxifen (OHT) induced *Cdc73* deletion in these cells (Fig. S4A) and impaired cell proliferation (Fig. 3A). In contrast, OHT treatment had no effect on control *CreER^T2^* cells. Next, we performed Bru-seq (Paulsen et al. 2013; Paulsen et al. 2014) in these cells to measure nascent mRNA transcripts following *Cdc73* deletion. We identified 1062 differentially expressed genes (“Cdc73 target genes”) shared between the two *Cdc73^f/f^* T-ALL cells but not by control cells (Fig. 3B, S4B). Consistent with previous studies, we did not observe large scale downregulation of gene of expression, which indicates that Cdc73 regulates specific genes.

**Figure 3.**
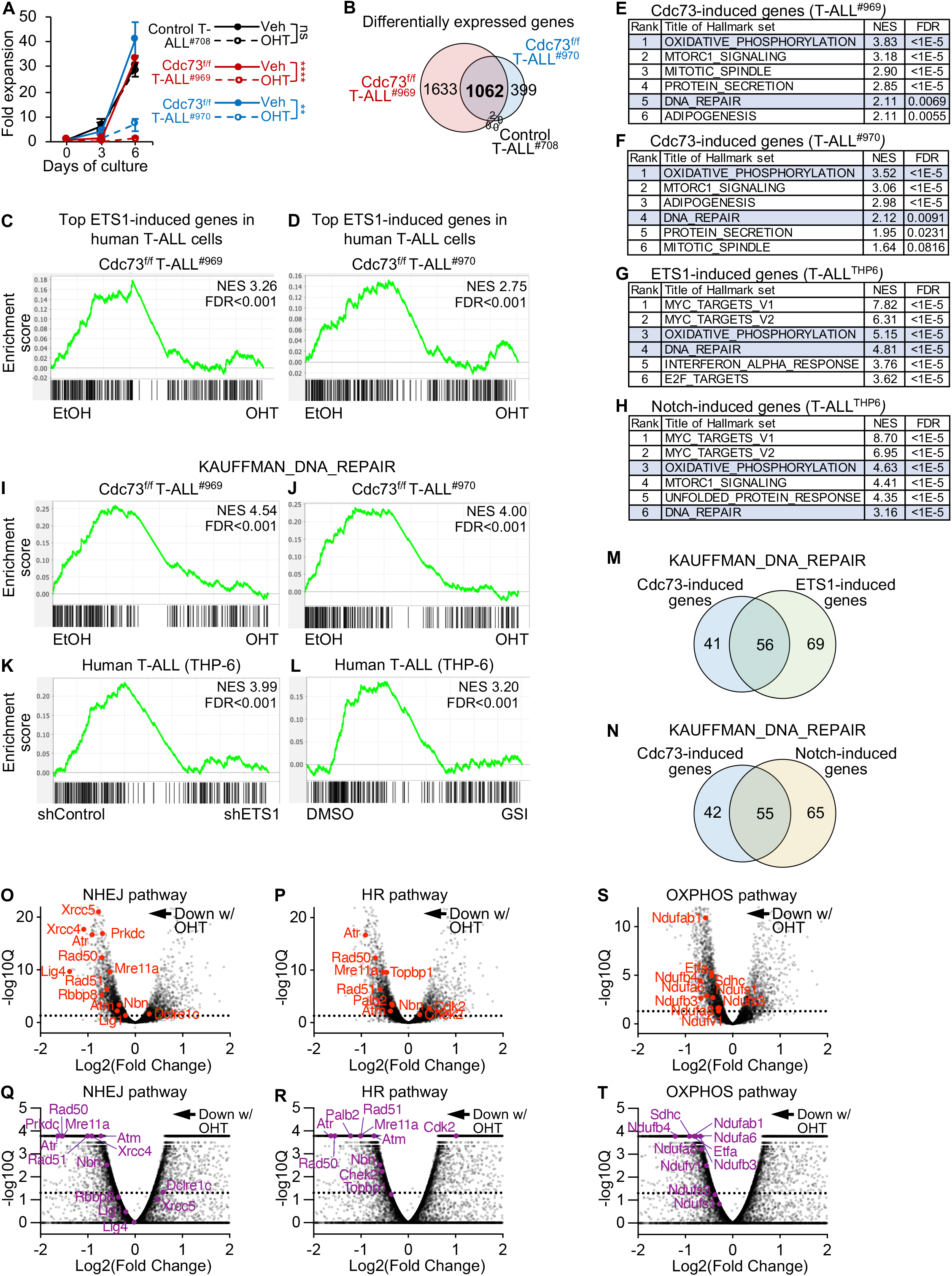
Cdc73 shares ETS1 and Notch-driven pathways. A) *Rosa26CreER^T2^ Cdc73^f/f^* T-ALL cells (969; 970) and control *Rosa26CreER^T2^* T-ALL cells derived from tumors in Fig. 2C were treated with 3nM OHT (hydroxytamoxifen) to delete *Cdc73* and measured for growth. Fold expansion=Day3 or Day6 cell count/Day 0 cell count. B) Venn diagram showing 1062 Cdc73 target genes shared by both *Cdc73^f/f^* T-ALL cell lines from (A). Target genes defined as FC>1.5; Padj<0.05 in Bru-Seq counts at 30h after OHT addition in both *Cdc73^f/f^* cell lines but not in controls. C-D) GSEA using the top 258 ETS1-induced gene list (McCarter et al., 2020) of Cdc73-induced genes in 969 (C) and 970 (D) cells. E-H) GSEA analyses showing the top 6 Hallmark pathways enriched for Cdc73-(E-F), ETS1-(G) and Notch-(H) induced target genes. ETS1 and Notch target genes were previously described in human THP-6 T-ALL cells (McCarter et al., 2020). Pathways shared across the 4 analyses are highlighted in blue. I-L) GSEA using the MSigDB C2 Kauffman_DNA_repair gene list of Cdc73-induced genes (I-J), ETS1-induced genes (K), and Notch-induced genes (L). M-N) Venn diagram showing overlap of Cdc73-induced and ETS1-induced (M) or Notch-induced (N) genes in the Kauffman_DNA_repair gene list. O-T) Volcano plots of significance vs. Bru-Seq (O-P, S) or RNA-Seq (Q-R, T) Log2FC data showing the control vs. OHT (Cdc73^Δ/Δ^) comparison and highlighting important genes in non-homologous end-joining (NHEJ; O, Q), homologous recombination (HR; P, R), and oxidative phosphorylation (OXPHOS; S-T) pathways in 969 T-ALL cells (O-P, S) and AML cells (Q-R, T) on background of all genes (grey) giving average RPKM>0.8. RNA-Seq analysis of Control and *Cdc73*-deleted AML cells were obtained from (Saha et al. 2019). ^**^P<0.01; ^****^P<0.0001.

Gene set enrichment analysis (GSEA) using a list of high-confidence direct NOTCH1 target genes in T-ALL (Wang et al. 2014) showed no significant enrichment for Cdc73-regulated target genes (Fig. S4C-D). However, *Cdc73* deletion significantly impaired the expression of 25% of these genes in both *Cdc73^f/f^* T-ALL cell lines at q<0.05 (P=0.0219, Fisher exact test). In contrast, GSEA using a list of high-confidence direct ETS1 target genes in T-ALL (McCarter et al. 2020) showed strongly significant enrichment for Cdc73-regulated genes in both 969 cells (*Cdc73^f/f^* T-ALL cell line; NES 3.26; FDR<0.001) and 970 cells (*Cdc73^f/f^* T-ALL cell line; NES 2.75; FDR<0.001) (Fig. 3C-D). *Cdc73* deletion impaired expression of 22% of ETS1 signature genes at q<0.05 (P=0.0003, Fisher exact test). These data are consistent with our protein-protein interaction and ChIP-Seq data showing stronger interaction of CDC73 with ETS1 than with NOTCH1 (Fig. S1A-C, S1F-G). Hallmark GSEA analysis identified DNA repair and oxidative phosphorylation (OXPHOS) gene signatures as the top two shared pathways enriched for Cdc73-induced, ETS1-induced, and Notch-induced genes (highlighted in blue in Fig. 3E-H). Thus, Cdc73 shares essential roles for T-cell development and leukemogenesis with Notch1 and Ets1 but has only partially overlapping functions with these factors in regulating gene expression.

DNA repair might be a synthetic lethal vulnerability in T-ALL due to oncogenic stress (Kotsantis et al. 2018; Leon et al. 2020). The most enriched DNA repair signature for Cdc73-induced genes in MSigDB was Kauffman_DNA_repair_genes, which gave FDR<0.001 for Cdc73-induced genes (NES 4-4.54, Fig. 3I-J), ETS1-induced genes (NES 3.94, Fig. 3K), and Notch-induced genes (NES 3.2, Fig. 3L). Analysis of core enrichment genes showed that ETS1 and Notch1 induce large fractions of Cdc73-induced DNA repair genes (58% in Fig. 3M and 58% in Fig. 3N respectively). Conversely, Cdc73 induces large fractions of ETS1 and Notch-induced DNA repair genes (45% and 46% respectively). *Cdc73* deletion generally downregulated important DNA repair genes in multiple pathways, such as non-homologous end joining (NHEJ; Fig. 3O, S4E) and homologous recombination (HR; Fig. 3P, S4F). To test if Cdc73 functions apply to Notch-independent and non-lymphoid contexts, we interrogated our gene expression datasets of murine MLL-AF9 driven *Cdc73^f/f^* acute myeloid leukemia (AML) cells (Saha et al. 2019). Accordingly, *Cdc73* deletion in these cells led to general downregulation of the same DNA repair genes (Fig. 3Q-R).

OXPHOS is a synthetic lethal vulnerability in T-ALL in part because Notch promotes high anabolic demand (Palomero et al. 2006; Kishton et al. 2016; Garcia-Bermudez et al. 2018; da Silva-Diz et al. 2021; Thandapani et al. 2021; Baran et al. 2022). Of the OXPHOS gene lists in MSigDB, the Hallmark list gave the highest enrichment with FDR<0.001 for Cdc73-induced genes (NES 3.52-3.83, Fig. S4G-H), ETS1-induced genes (NES 5.15, Fig. S4I), and Notch-induced genes (NES 4.63, Fig. S4J). Analysis of core enrichment genes showed that ETS1 and Notch induce large fractions of Cdc73-induced OXPHOS genes (60% in Fig. S4K and 62% in Fig. S4L respectively). Conversely, Cdc73 induces large fractions of ETS1 and Notch-induced OXPHOS genes (56% and 41% respectively). *Cdc73* deletion generally downregulated important OXPHOS pathway genes (Fig. 3S, S4M). These same genes were also generally downregulated upon *Cdc73* deletion in AML cells (Fig. 3T) (Saha et al. 2019). Expression analysis of sorted DN subsets showed loss of important DNA repair and OXPHOS genes, particularly at the DN4 cell stage when the strongest Cdc73 deletion was achieved (Fig. S4N). Taken together, like ETS1 and Notch, Cdc73 promotes expression of genes that are important for DNA repair and OXPHOS. These roles appear to be conserved among early T cells, T-ALL cells, and Notch-independent/non-lymphoid AML cells.

### Cdc73 is important for genome integrity

Previous work by others showed that CDC73 is important for genome stability through maintenance of telomere length, homologous recombination repair, transcription-coupled repair, and R-loop clearance at sites of transcriptional elongation (Gaillard et al. 2009; Tatum et al. 2011; Wahba et al. 2011; Herr et al. 2015; Nene et al. 2018; Shivji et al. 2018). DepMap analysis of several cancer types showed that T-ALL cells express the highest levels of DNA repair target genes that are shared between Cdc73, Notch and Ets1 (leftmost group in Fig. S5A). Given these data, we wondered whether Cdc73 helps preserve genome integrity in T-ALL. To test this, we treated *Cdc73^f/f^* T-ALL cell lines with OHT and measured levels of γH2AX, a marker of DNA double-strand breaks. Consistently, OHT treatment of both *Cdc73^f/f^* T-ALL cell lines, but not the control cell line, effectively suppressed Cdc73 expression and increased γH2AX levels (Fig. 4A). Further, CDC73 knockdown had similar effects in SUP-T1 and CUTLL1, two human T-ALL cell lines that, like the *Cdc73^f/f^* T-ALL cell lines, are Notch-induced (Fig. 4B). Since γH2AX signals cluster at promoters of genes regardless of expression level in T-ALL cell lines (Seo et al. 2012), we examined localization of γH2AX at the promoters of a panel of genes. Consistently, qChIP showed that *Cdc73* deletion induced γH2AX localization to chromatin (Fig. S5B). In contrast, OHT had no effect in control cells (Fig. S5C). To confirm that *Cdc73* deletion led to spontaneous DNA damage, we performed blinded assays to assess chromosomal damage in metaphase spreads. Consistently, we observed increased chromosomal abnormalities in Cdc73^f/f^ T-ALL metaphases upon treatment with OHT (Fig. 4C-E). As expected, DNA damage upon OHT treatment led to a small but significant increase in apoptosis (~4-fold, Fig. 4F-G). Next, we wondered whether T-ALL cells are sensitive to DNA repair inhibition. To test this possibility, we treated *Cdc73^f/f^* and human T-ALL cells with berzosertib, an inhibitor of the apical DNA damage protein kinase, ATR, which is being tested in advanced clinical trials due to its excellent safety profile. Consistently, these cells were highly sensitive to ATR inhibition given low nanomolar GI50 (Fig. 4H-I). Dependency on DNA repair is shared by cancers in general as non-hematopoietic cancer cell lines were also sensitive and ATR labeled as a “common essential” gene in DepMap 22Q4. These data suggest that Cdc73 promotes DNA repair globally through gene expression, which extends previous studies and might help protect T-ALL cells from chromosomal damage.

**Figure 4.**
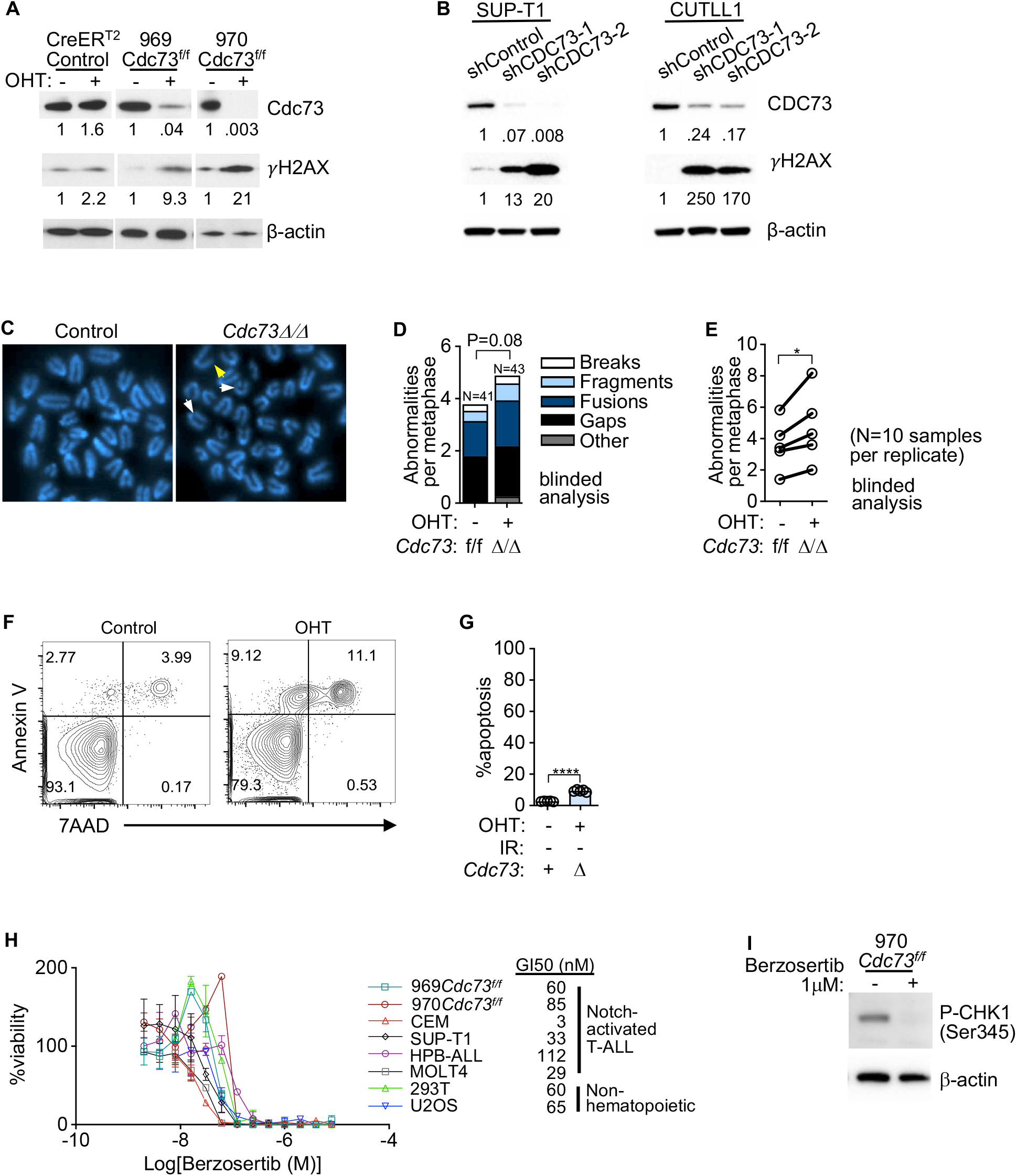
Cdc73 is important for genome integrity. A) Western blot for *γ*H2AX in *Rosa26CreER^T2^ Cdc73^f/f^* T-ALL cells (969; 970) and control *Rosa26CreER^T2^* T-ALL cells treated with OHT (hydroxytamoxifen) for 30 hours to delete *Cdc73*. Numbers represent band intensities normalized to β-actin loading control. B) Western blot for *γ*H2AX in human NOTCH1-induced human T-ALL cell lines (SUP-T1; CUTLL1) transduced with shCDC73. C-E) Representative metaphase spreads (C) and quantification of metaphase abnormalities in aggregate (D) or per replicate (E) in blinded analyses of *Cdc73^f/f^* T-ALL cells treated with OHT for 30 hours. White arrows=gaps; yellow arrow=break. F-G) Representative Annexin V/7-AAD flow cytometric plots (F) and Annexin V^+^/7-AAD^−^ scatterplot (G) of *Cdc73^f/f^* T-ALL cells treated with vehicle (Control) or OHT for 30 hours (*Cdc73Δ/Δ*). H) Dose response curves of Notch-activated T-ALL cell lines and 2 non-hematopoietic tumor cell lines (U2OS and 293T) treated with berzosertib (ATR inhibitor in clinical trials). I) Western blot in UV-treated 970 cells showing effect of berzosertib on P-Chk1, a downstream target of Atr.

### Cdc73 is important for oxidative phosphorylation

Supraphysiological Notch signals in T-ALL have been previously shown to directly and indirectly induce OXPHOS genes, such as electron transport Complex I (Kishton et al. 2016; Baran et al. 2022). DepMap analysis of several cancer types showed that T-ALL cells express average levels of OXPHOS target genes that are shared between Cdc73, Notch and Ets1 (leftmost group in Fig. S6). Since Cdc73 promotes expression of genes important for the electron transport chain (Fig. 3S, S4M), we considered the possibility that Cdc73 helps maintain membrane potential to protect T-ALL cells from metabolic stress. To test this possibility, we treated *Cdc73^f/f^* T-ALL cell lines with OHT and measured mitochondrial membrane potential of live cells using the tetramethylrhodamine methyl ester assay (TMRM) (Fig. 5A-B). *Cdc73* deletion reduced membrane potential to nearly the same degree as FCCP, a mitochondrial uncoupler. Based on our gene expression analysis showing that Cdc73 regulates Complex I (Fig. 3S, S4M), we predicted that *Cdc73* deletion would reduce reactive oxygen species (ROS). Consistently, the

**Figure 5.**
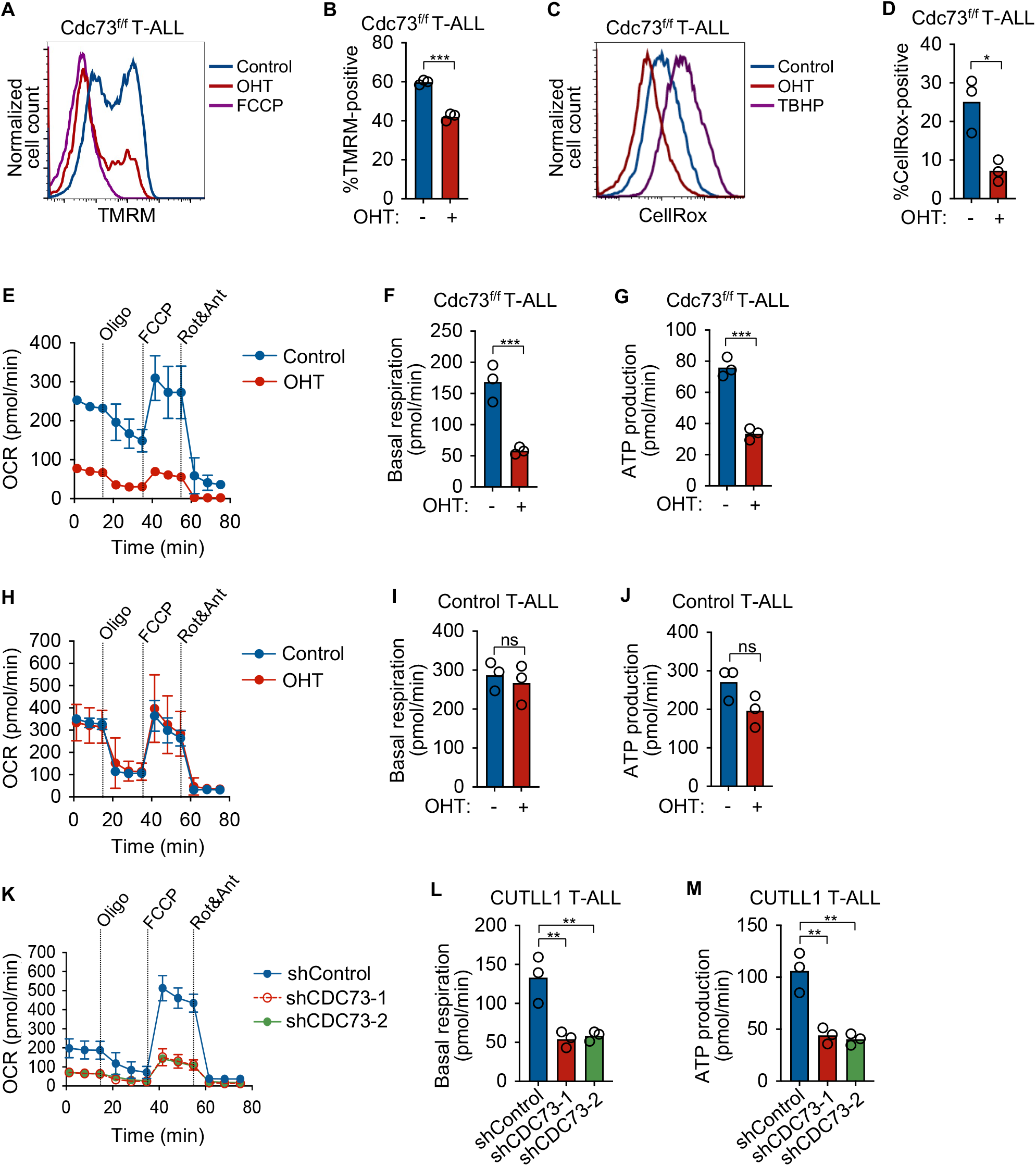
Cdc73 is important for oxidative phosphorylation. A-B) Representative flow cytometric histogram (A) and MFI scatterplot (N=3) (B) showing mitochondrial membrane potentials measured by tetramethylrhodamine methyl ester (TMRM) assay on live Dapi-*Rosa26CreER^T2^ Cdc73^f/f^* T-ALL cells (970) treated with OHT (hydroxytamoxifen) for 30 hours to delete *Cdc73*. FCCP was added as positive control. C-D) Representative flow cytometric histogram (C) and MFI scatterplot (N=3) (D) showing mitochondrial ROS production measured by the CellROX assay on live Dapi-970 cells treated with OHT for 30 hours to delete *Cdc73*. TBHP was added as a positive control. E-M) Seahorse XFe96 instrument measurements of real time oxygen consumption rate (OCR) normalized to live cell number and protein concentration under basal conditions or in response to the indicated mitochondrial inhibitors (E,H,K) and scatterplots of basal (F,I,L) and ATP production (G,J,M) respiration phases of 970 cells treated with OHT for 30 hours to delete *Cdc73* (E-G), control *Rosa26CreER^T2^* cells treated with OHT for 30 hours (H-J); and CUTLL1 cells at 4 days after transduction with two independent shCDC73 (K-M).

CellROX assay showed that OHT treatment reduced ROS production (Fig. 5C-D). Our gene expression analyses also predicted impaired oxygen consumption. Consistently, the mitochondrial stress test showed that *Cdc73* deletion reduced basal respiration and mitochondrial ATP production rates of live cells (Fig. 5E-G). In contrast, treating control T-ALL cells with OHT had no significant effect (Fig. 5H-J). CDC73 knockdown in live human Notch-induced T-ALL cells also impaired oxygen consumption rates (Fig. 5K-M). These data suggest that Cdc73 promotes OXPHOS, which might help protect T-ALL cells from the metabolic stresses of supraphysiological Notch signaling.

### Cdc73 does not primarily promote DNA repair and OXPHOS gene expression through enhancers

Previous groups showed that Paf1C regulates enhancer activity by altering eRNA synthesis and H3K27ac deposition (Chen et al. 2017; Ding et al. 2021). Consistently, we observed strong CDC73 ChIP-Seq signals at intergenic regulatory elements relative to promoters (Fig. S7A). To test whether Cdc73 regulates enhancer activity in T-ALL, we performed H3K27ac ChIP-Seq in our *CreER^T2^ Cdc73^f/f^* T-ALL cell lines (969 and 970) and control *CreER^T2^* T-ALL cell line after OHT treatment. In support of enhancer functions for Cdc73, ChIP-Seq showed differential H3K27ac signals upon *Cdc73* deletion at FDR<0.05 (Fig. 6A-B, S7B). In contrast, OHT had little effect on control cells (Fig. 6C). We intersected the datasets to obtain 9,139 “dynamic H3K27ac peaks”, which were defined based on differential H3K27ac signals at FDR<0.05 in the same direction for both *CreER^T2^ Cdc73^f/f^* T-ALL cell lines but not the control *CreER^T2^* T-ALL cell line. In general, *Cdc73* deletion impaired H3K27ac signals at dynamic intergenic H3K27ac peaks (Fig. 6D, S7C). These data suggest that Cdc73 promotes enhancer activity more than it represses enhancer activity in T-ALL.

**Figure 6.**
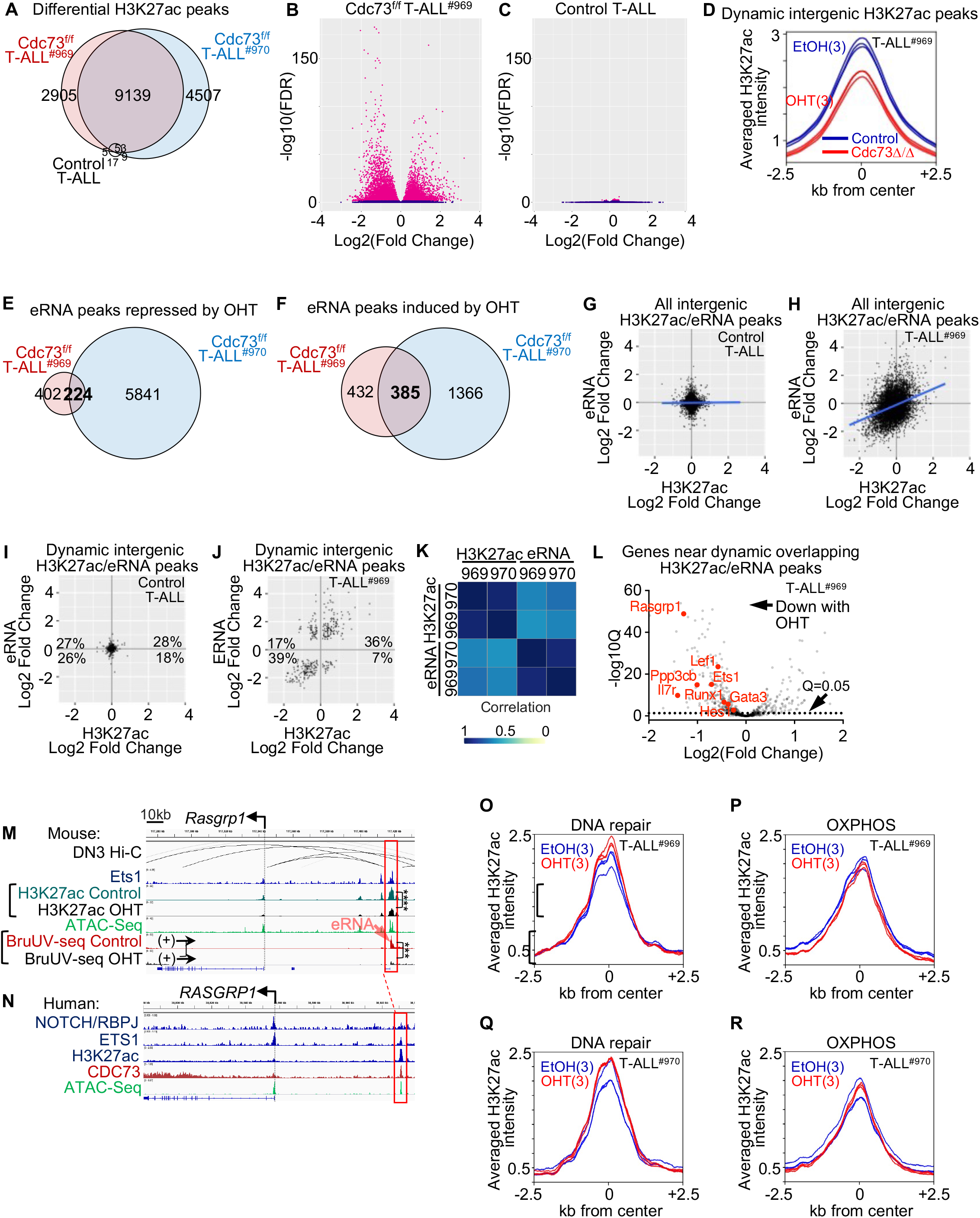
Cdc73 does not primarily promote DNA repair and oxidative phosphorylation gene expression through enhancers. A) Venn diagram of differential H3K27ac (FDR<0.05) in *Rosa26CreER^T2^ Cdc73^f/f^* T-ALL cells (969; 970) and control *Rosa26CreER^T2^* T-ALL cells upon treatment with 6nM OHT for 30 hours. B-C) Volcano plots of significance vs. Log2(OHT/Control) H3K27ac ChIP-Seq signals of 969 (B) and control (C) cells in (A). D) Metagene plot of dynamic intergenic H3K27ac signals (defined as FDR<0.05 in 969 and 970 cells but not in control cells) in 969 cells. E-F) Venn diagram of differential eRNAs in *Cdc73^f/f^* T-ALL cells (969; 970) that were repressed (E) or induced (F) upon treatment with 6nM OHT for 30 hours. eRNAs were defined as intergenic BruUV-Seq peaks or intragenic peaks that were antisense in direction relative to mRNAs. No differential eRNAs were identified after OHT treatment of control T-ALL cells. G-J) BruUV-Seq Log2(OHT/Control) versus H3K27ac Log2(OHT/Control) scatterplots of all overlapping intergenic peaks (G-H) or overlapping dynamic intergenic peaks (I-J) in control T-ALL cells (G, I) and *Cdc73^f/f^* T-ALL cells (969; H, J). Overlapping “dynamic peaks” were defined as giving q<0.05 and FDR<0.05 in the same direction for the BruUV-Seq and H3K27ac comparisons respectively in both *Cdc73^f/f^* T-ALL cells but not in control T-ALL cells. K) Spearman’s correlation coefficient analysis of eRNA and H3K27ac Log2(OHT/Control) from (J and Fig. S7G) in 969 and 970 *Cdc73^f/f^* T-ALL cells. L) Volcano plot of significance vs. Bru-Seq Log2(OHT/Control) of genes nearest overlapping OHT-downregulated dynamic intergenic BruUV-Seq and H3K27ac peaks in 969 *Cdc73^f/f^* T-ALL cells. M-N) Display tracks of indicated ChIP-Seq and ATAC-Seq datasets at the *Rasgrp1* locus in mouse 969 cells (M) or human THP-6 cells (N) showing nearest mouse-human homologous enhancers in red boxes that contain overlapping dynamic intergenic eRNA and H3K27ac peaks. Ets1 ChIP-seq (GSM461516); ATAC-seq (GSM2461649); DN3 Hi-C (GSE79422) analyzed in (Kashiwagi et al. 2022). O-R) Metagene plots of H3K27ac signals at non-promoter H2K27ac peaks nearest DNA repair (O, Q) and OXPHOS genes (P, R) from Table S1 in 969 cells (O-P) and 970 cells (Q-R). ^**^FDR<0.01; ^***^FDR<0.001; NS=not significant.

Since Paf1C regulates enhancer activity through eRNA synthesis (Chen et al. 2017; Ding et al. 2021), we next sought to identify enhancers directly regulated by Cdc73 by integrating our dynamic H3K27ac dataset with differential analysis of eRNA expression. Towards this goal, we performed BruUV-seq on the *CreER^T2^ Cdc73^f/f^* T-ALL cell lines (969 and 970) and control *CreER^T2^* T-ALL cell line. BruUV-seq is a nascent RNA technique that detects and quantifies rapidly degraded RNAs like eRNAs and is performed using living cells (Magnuson et al. 2015). We identified 609 differential eRNAs at q<0.05 upon *Cdc73* deletion that were shared by both *Cdc73^f/f^* T-ALL cell lines but not by the control cell line (Fig. 6E-F, S7D-E). Next, we integrated the H3K27ac and BruUV-seq datasets to identify intergenic H3K27ac and eRNA peaks that overlapped in both T-ALL cell lines. There was no correlation between changes in eRNA and H3K27ac signal in control cells (Fig. 6G). In contrast, there was modest correlation in *Cdc73^f/f^* T-ALL cells (Fig. 6H, S7F). To determine the genomic locations where Cdc73 might have strong direct effects on enhancers, we defined “dynamic eRNA peaks” as giving q<0.05 in the same direction for the BruUV-seq comparison in both *Cdc73^f/f^* T-ALL cells but not in control T-ALL cells. Changes in dynamic intergenic H3K27ac signals showed no correlation in control cells (Fig. 6I) but strong correlation in both *Cdc73^f/f^* T-ALL cells with changes in dynamic intergenic eRNA signals (r>0.66; Fig. 6J-K, S7G). Our data now extends to T-ALL previous observations that CDC73 coordinately regulates eRNA synthesis and enhancer activity.

Next, we sought to identify direct target genes of Cdc73 by associating overlapping dynamic intergenic H3K27ac/eRNA peaks that diminish upon *Cdc73* deletion with nearest expressed genes. We found that OHT-downregulated H3K27ac/eRNA peaks were associated with OHT-downregulated expression of several known genes that are important for maintaining T-ALL proliferation, such as *Il7r, Ets1*, *Rasgrp1*, and *Lef1* (Hartzell et al. 2013; Ksionda et al. 2016; Oliveira et al. 2019; Silva et al. 2021; Carr et al. 2022) (Fig. 6L-N, S7H-N). *Il7r* and *Lef1* are known direct Notch target genes (Spaulding et al. 2007; Gonzalez-Garcia et al. 2009; Wang et al. 2014).

Importantly, we consistently observed CDC73 occupancy at homologous human enhancers (Fig. 6N, S7J, S7L, S7N). RBPJ/Notch and ETS1 occupancy was often but not consistently observed. Analysis of publicly available Hi-C datasets of DN3 cells confirmed interactions of these enhancers with respective promoters, except for the previously described Notch-dependent *Il7r* enhancer (Wang et al. 2014), presumably because DN3 cells express little *Il7r*. Interestingly, we did not observe association of overlapping dynamic intergenic H3K27ac/eRNA peaks with core enrichment genes from GSEA pathway analyses in DNA repair (Fig. 3I-J, Table S1) or OXPHOS (Fig. S4G-H, Table S1). Furthermore, we did not observe any differences in H3K27ac signal changes upon *Cdc73* deletion at non-promoter peaks nearest DNA repair or OXPHOS genes relative to all genes in both cell lines (Fig. 6O-R, Fig. S7O-P). These data suggest that Cdc73 activates enhancers that induce important T-ALL driver genes. However, these non-canonical functions do not appear to induce expression of DNA repair and OXPHOS genes.

### Cdc73 promotes DNA repair and OXPHOS gene expression through canonical functions at gene bodies

Since Cdc73 did not appear to induce expression of DNA repair and OXPHOS genes through enhancers, we considered the possibility that Cdc73 induces these genes through its well-established functions in promoting mRNA synthesis. To test this, we first counted CDC73 tags between the transcriptional start site (TSS) and transcriptional termination site (TTS) of GSEA core enrichment genes in DNA repair or OXPHOS (Table S1, Fig. 7A). Consistently, CDC73 tags were generally more abundant at DNA repair and OXPHOS genes relative to all genes. ETS1 and RBPJ/Notch tags were also more abundant at promoters of these genes (Fig. 7B-C). This observation is consistent with our ChIP-Seq and gene expression analyses showing that CDC73 intersects with the ETS1 and Notch pathways. Next, we performed H2BK120ub1 ChIP-Seq on our *CreER^T2^ Cdc73^f/f^* T-ALL cell lines upon treatment with OHT to delete *Cdc73*. We chose H2BK120ub1 since one of the best-established mRNA functions of Paf1C is to promote transcriptional elongation by recruiting the Bre1-Rad6 E3 ubiquitin ligase complex to monoubiquitinate H2BK120 (Ng et al. 2003; Kim et al. 2009; Kim and Roeder 2009; Van Oss et al. 2016; Chen et al. 2021). Further, Bre1-mutated flies show notched wings and defective Notch target gene expression (Bray et al. 2005).

**Figure 7.**
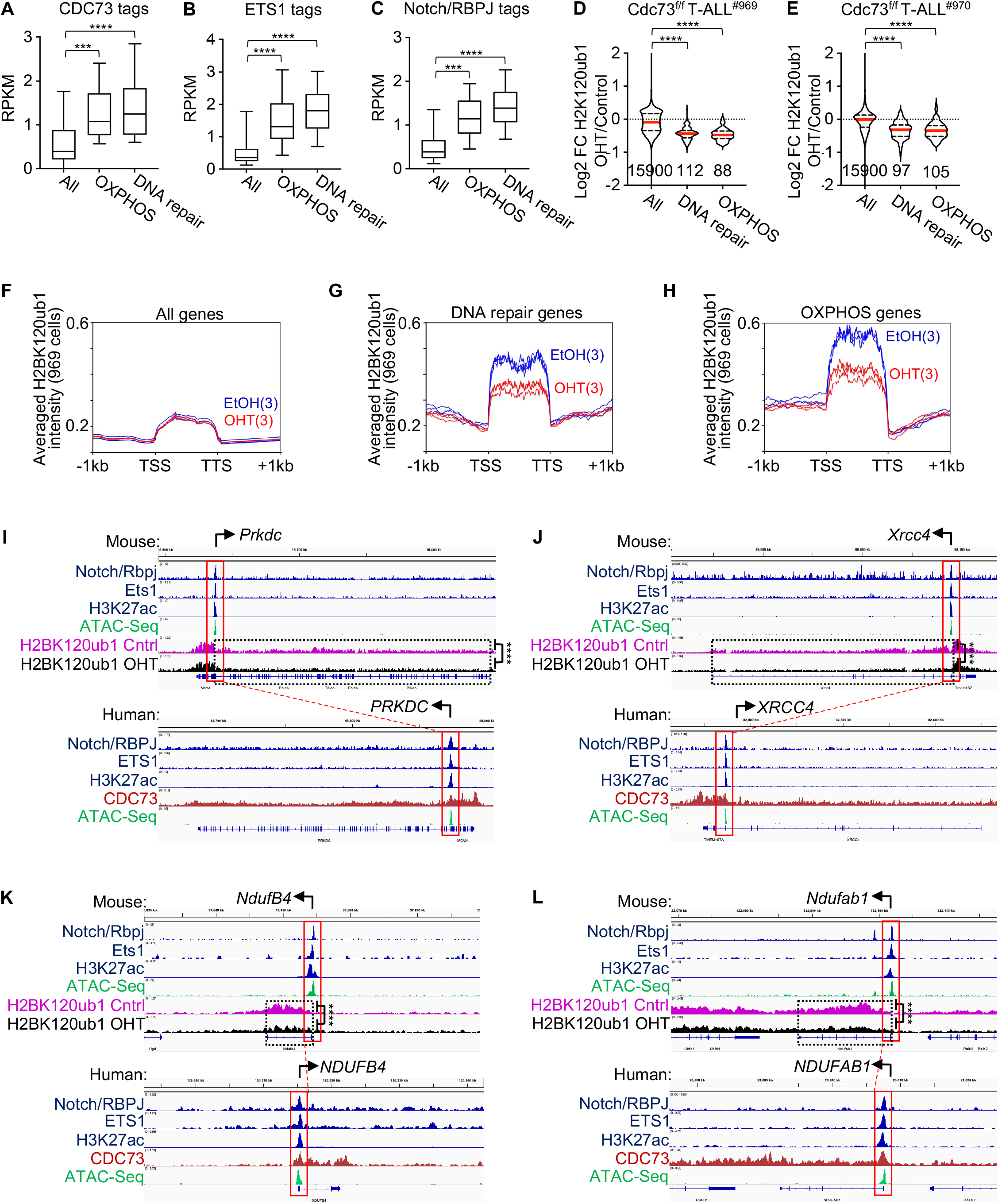
Cdc73 promotes DNA repair and OXPHOS gene expression through canonical mRNA functions at gene bodies. A-C) Box and whisker plots of CDC73 (A), ETS1 (B), and Notch/RBPJ (C) tag counts in human THP-6 cells at gene bodies (A) and promoters (B-C) of all genes, OXPHOS genes, and DNA repair genes shared by the GSEA enrichment cores in *Cdc73^f/f^* 969 and 970 T-ALL cells (Fig. 3I, Fig. S4G, Table S1). D-E). Violin plots showing H2BK120ub1 ChIP-Seq Log2FC in 969 cells (D) and 970 cells (E) for all genes, DNA repair genes, and OXPHOS genes in the GSEA enrichment cores of 969 and 970 cells (Table S1). OHT was added for 30 hours to delete *Cdc73*. F-H) Metagene plots of H2BK120ub1 signals in EtOH- (blue) and OHT- (red) treated 969 cells at all genes (F), DNA repair genes (G), and OXPHOS genes (H) in core enrichment genes of GSEA analyses (Table S1). I-L) Display tracks of indicated ChIP-Seq and ATAC-Seq datasets at important DNA repair genes (I-J) and OXPHOS genes (K-L) in mouse 969 cells (top) or human THP-6 cells (bottom) showing representative tracks and FDR values upon OHT addition (*Cdc73* deletion) of H2BK120ub1 signals between TSS and TTS. ^****^FDR<0.0001. ATAC-seq (GSM2461649). Ets1 ChIP-Seq (GSM2461515).

Consistent with its H2BK120 monoubiquitination functions, *Cdc73* deletion reduced the most H2BK120ub1 signal at mouse genes ranked in the top tercile of CDC73 signal at homologous human genes (Fig. S8A, S8D) relative to the middle tercile (Fig. S8B, S8E) and bottom tercile (Fig. S8C, S8F) in both *Cdc73^f/f^* cell lines. *Cdc73* deletion generally caused downregulation of H2BK120ub1 signals at DNA repair and OXPHOS genes relative to all genes in both *Cdc73^f/f^* cell lines (Fig. 7D-H, Fig. S8G-I). For example, we observed significant downregulation (FDR<0.01) of H2BK120ub1 counts at important DNA repair genes like *Prkdc*, *Xrcc4*, *Rad50* and *Atr* (Fig. 7I-J; Fig. S8J-K) and important OXPHOS genes like *NdufB4*, *Ndufab1*, *Etfa*, and *Ndufa6* (Fig. 7K-L, S8L-M). Importantly, broad CDC73 signals were consistently observed across gene bodies of the homologous human genes (bottom panels of Fig. 7I-L and Fig. S8J-M). As expected, based on our genome-wide analyses of ETS1 and RBPJ/Notch binding in Fig. 7B-C, we consistently observed occupancy of all these factors at the promoters of the mouse genes and human homologs. Core enrichment analysis showed that Notch induces 5 out the 8 genes while ETS1 induces all 8 genes. Our data suggest that CDC73 might intersect NOTCH1 and ETS1 occupancy at DNA repair and OXPHOS genes and directly induce mRNA synthesis of through promoting H2BK120 monoubiquitination.

## Discussion

Notch signaling controls developmentally important activities in diverse tissues, raising barriers for developing anti-Notch therapies that directly target Notch. Emerging evidence originating in *Drosophila* highlights an alternative strategy of targeting transcriptional regulators that co-bind Notch-occupied regulatory elements (Bray 2016; Falo-Sanjuan and Bray 2019). Previous work showed that NOTCH1 bound CDC73 in breast cancer cells and relied on Paf1C to induce Notch target gene expression for *Drosophila* wing development (Bray et al. 2005; Tenney et al. 2006; Kikuchi et al. 2016). Here, we confirmed the NOTCH1-CDC73 interaction in T-ALL cells. However, the interaction was context dependent. RBPJ/Notch co-occupied only a subset of CDC73-bound elements, which were highly enriched for ETS motifs and ETS1 occupancy. Further, ETS1 interacted with CDC73 and ETS1 knockdown generally impaired CDC73 binding to chromatin where ETS1 was also bound. Thus, we extend previous studies by showing that chromatin context (here ETS1 in T-ALL) might restrict the NOTCH1-CDC73 interaction.

Functionally, Ccd73 acts as a tumor suppressor in some cancers (Carpten et al. 2002; Wang et al. 2005; Hanks et al. 2014) and as an oncogene in others like AML (Muntean et al. 2010; Zeng and Xu 2015; Zhi et al. 2015; Karmakar et al. 2017; Saha et al. 2019), Here, we showed that the role of Cdc73 in T-ALL is consistent with the latter group. Accordingly, *Cdc73* deletion in mice impaired Notch-dependent T-cell development and leukemogenesis in mouse and human models. Consistent with our chromatin profiling, gene expression analyses showed that Notch and Ets1 converge only on a subset of Cdc73-induced pathways, notably boosting ~60% of Cdc73 target genes in DNA repair and OXPHOS through canonical functions and key oncogenes like *Il7r* and *Lef1* through enhancer functions. Conversely, CDC73 intersects and supports a subset of Notch and ETS1 functions. We show no evidence that Cdc73 performs these actions exclusively in the Notch-dependent or T-ALL-specific context. On the contrary, Cdc73 appears to have similar functions in MLL-driven AML cells, which are Notch-independent and non-lymphoid.

Previous work by others showed that CDC73 is important for genome stability at telomeres and actively transcribed genes (Gaillard et al. 2009; Tatum et al. 2011; Wahba et al. 2011; Herr et al. 2015; Nene et al. 2018; Shivji et al. 2018). Here, we extend this work by showing that Cdc73 might have more global effects by promoting expression of DNA repair genes, particularly those required for double-stranded DNA break repair, and often intersecting with the Notch pathway at ~60% of these genes. Consistently, *Cdc73* deletion increased chromosomal damage and induced *γ*H2AX. Previous groups showed that T-ALL cells are highly dependent on the electron transport chain since Notch-dependent and independent pathways upregulate MYC and mTORC1, raising demand for ATP to fuel anabolic processes (Palomero et al. 2006; Kishton et al. 2016; Garcia-Bermudez et al. 2018; da Silva-Diz et al. 2021; Thandapani et al. 2021). Here, we show that Cdc73 might help replenish cellular energy by boosting OXPHOS target genes, often intersecting with the Notch pathway at ~60% of these genes. Together with previous studies, our data suggest that Cdc73 promoted a gene expression program important for DNA repair and mitochondrial function. At some point, Notch intersected with Cdc73 at these genes to mitigate the genotoxic and metabolic stresses of elevated Notch signaling (Fig. S9).

Previous studies by others showed divergent roles of the Paf1C in regulating enhancer activity in colon cancer cells and embryonic stem cells (Chen et al. 2017; Ding et al. 2021). In the colon cancer study (Chen et al. 2017), PAF1 repressed eRNA transcription and enhancer activity by maintaining the paused state of Pol II (Chen et al. 2015). In contrast, in the ES cell study (Ding et al. 2021), Ctr9 stimulated eRNA transcription and enhancer activity, consistent with the consensus view of Paf1C as an activator of transcription (Jaehning 2010; Yu et al. 2015). These complexities might be explained by context dependence or by another positive function for Paf1C in promoting transcriptional elongation beyond the promoter (Hou et al. 2019). Consistent with the consensus view of Paf1C, our integrated analysis showed that *Cdc73* deletion impaired H3K27ac signals and eRNA synthesis at CDC73-bound enhancers associated with known T-ALL drivers. However, differential enhancer activity was not linked to differential expression of Cdc73-induced DNA repair or OXPHOS genes. Instead, integrated analysis of CDC73 and H2BK120ub1 signals suggested that Cdc73 primarily induces these genes by promoting mRNA synthesis, which is the consensus primary function of Paf1C. Our interpretation is in line with the consensus view that enhancers regulate lineage and developmental stage-specific gene expression while promoters regulate housekeeping gene expression.

Therapeutically, our investigation suggests that DNA repair, oxidative phosphorylation, and CDC73 might be therapeutic targets in T-ALL. However, we are mindful that targeting CDC73 might be toxic based on studies of ubiquitous *Cdc73* deletion (Wang et al. 2008). Thus, future studies on the CDC73 interactome are needed to find ways to selectively target CDC73 functions, such as through inhibitors of protein-protein interactions. Further, our study supports the notion that T-ALL cells are a cancer type that is sensitive to inhibition of general transcriptional machinery, which had once been considered to be too toxic to block given its vital functions in normal cells (Ott 2014). Previous work showed that T-ALL cells are highly dependent on cofactors of Pol II, such as CDK7, MYC, and BRD4 (Bisgrove et al. 2007; Rahl et al. 2010; Kwiatkowski et al. 2014; Roderick et al. 2014). Here we extend so-called “transcriptional addiction” in T-ALL to CDC73. Previously, we showed that leukemia-associated NOTCH1 alleles are weak transactivators (Chiang et al. 2008). Our current work suggests that oncogenic Notch might overcome its inherent weakness through local intersection at chromatin with co-binding transcription factors and powerful RNA synthesis machinery, which induce the most highly active chromatin and expressed genes. These intersecting pathways might therefore represent vulnerabilities that could be exploited to target Notch signals without directly targeting the Notch complex, thus mitigating toxicities.

## Materials and methods

### Mice

C57/BL6 mice between the ages of 4-8 weeks and *LckCre* mice were purchased from Taconic. *Rosa26CreER^T2^ Cdc73^f/f^* mice were generated by crossing *Cdc73^f/f^* mice (a gift from Andy Muntean) (Wang et al. 2008) with *Rosa26CreER^T^*^2^ mice (Jackson Labs). *Cdc73^f/f^* mice were backcrossed to C57/BL6 strain at least five times prior to use. All mouse experiments were performed according to NIH guidelines and approved protocols from the Institutional Animal Care and Use Committee at the University of Michigan (Ann Arbor).

### Cell lines

CUTLL1 cells were obtained from Adolfo Ferrando and Andrew Weng. All other cell lines including SUP-T1, LOUCY, HPB-ALL, CEM, THP-6, and OP9-DL4 cells were obtained as previously published (McCarter et al. 2020). All human cell lines were authenticated using STR analysis prior to use (Genetica Corporation). Primary mouse T-ALL cell lines were derived by harvesting splenic or thymic tumors from *Rosa26CreER^T2^ Cdc73^f/f^* or *Rosa26CreER^T2^ Cdc73^+/+^* mice and culturing in 20% RPMI until growing well and then in 15% RPMI thereafter. All cell lines were cultured less than 3 months after resuscitation and tested for contaminants using MycoAlert (Lonza) every 1-3 months to ensure they were free of *Mycoplasma* contamination.

### Antibodies and primers

These reagents are listed in Table S2.

### Cell culture conditions

T-ALL cell lines were grown in RPMI1640 (Invitrogen) supplemented with FBS (Gibco), 2 mM l-glutamine, 2-mercaptoethanol [0.0005% (v/v), Sigma], penicillin, and streptomycin. 293T cells were maintained in DMEM (Invitrogen) with the same supplements except 2-mercaptoethanol. All cells were grown at 37°C under 5% CO_2_. 4-hydroxytamoxifen was obtained from Sigma, diluted in ethanol, and used at 0.1% final concentration of ethanol. Puromycin (Sigma) was added to transduced cell cultures 48 hours after transduction for knockdown experiments.

### Constructs and viral production

Flag-ETS1 (McCarter et al. 2020), Flag-NOTCH1 (Pinnell et al. 2015), and Flag-CDC73 (Ropa et al. 2018) constructs were previously described. ShRNA constructs were obtained from Sigma: shControl (SHC002), shCDC73-1 (TRCN0000329742), and shCDC73-2 (TRCN0000329681). Retroviral and lentiviral production and titering was performed as previously described (McCarter et al. 2020). Lentivirus was concentrated using 30 kDa MWCO Amicon Ultra filters (Sigma).

### Human patient expression data

Human patient data was based upon data generated by Therapeutically Applicable Research to Generate Effective Treatments (TARGET; https://ocg.cancer.gov/programs/target) initiative, phs000218. The ALL project team was headed by Stephen P. Hunger, M.D. at the University of Colorado Cancer Center, Denver, CO, USA. The dbGaP Sub-study ID was phs000463/phs000464. The data used for this analysis are available at https://portal.gdc.cancer.gov/projects.

### PDX experiments

PDX3 (M71) and PDX4 (2583AB) were obtained from Andrew Weng and Moshe Talpaz respectively. PDXs were expanded by injection into nonirradiated NOD-*scid-*IL2γ^null^ (NSG) mice. De-identified human samples were obtained and used with approval from the University of Michigan Institutional Review Board and informed consent under guidelines established by the Declaration of Helsinki. Generation of PDX samples, transduction of PDX cells, and leukemia initiation experiments were performed as previously described (Yost et al. 2013; McCarter et al. 2020) with transplantation into nonirradiated NSG mice followed by injection into a second cohort of NSG mice for survival analysis after 24 weeks. Mice injected with PDX cells were sacrificed when moribund.

### Bone marrow transplantation

Bone marrow stem and progenitor cells of *Rosa26CreER^T2^* or *Rosa26CreER^T2^Cdc73^f/f^* mice were transduced with an activated *Notch1* allele (ΔE/Notch1). Transduced cells were injected into irradiated C57BL/6 mice to generate primary T-ALL tumors as previously described (Pear et al. 1996; Aster et al. 2000; McCarter et al. 2020).

### Flow cytometry

Flow cytometry and cell sorting was performed as previously described (McCarter et al. 2020). Mitochondrial potential staining was done using TMRM (ThermoFisher). FCCP was obtained from Sigma. Reactive oxygen species staining was done using CellRox DeepRed (ThermoFisher) and TBHP (Sigma). Each experimental condition was run in triplicate. The values displayed are representative of three biological replicates.

### Co-immunoprecipitation (co-IP) and western blot

Flag-tagged co-IP was performed as previously described (Pinnell et al. 2015). Endogenous co-IP and Western blot was performed as previously described (McCarter et al. 2020).

### Dose response curves

Human and murine T-ALL cells were treated at increasing doses of berzosertib (Chemietek) for 9-10 days. CEM and LOUCY cells were treated for 16 days. Cells were stained with AO/PI and then counted using a CellDrop cell counter (DeNovix).

### Quantitative PCR

QRT-PCR samples were prepared by extracting total RNA using the RNeasy Plus Mini Kit (Qiagen) according to manufacturer’s protocol. Random primed total RNA (0.5ug) was reverse transcribed with SuperScript IV (Invitrogen). TaqMan Universal PCR Master Mix or Power SYBR Green PCR Master Mix (Applied Biosciences) were used to amplify transcripts using QuantStudio3 Pro Real-Time PCR System (Applied Biosystems). Relative expression of target genes was measured comparing against housekeeping controls β-actin or Ef1a.

### Seahorse assay

Mitochondrial stress tests were performed using a Seahorse assay as described in a previously published protocol (Kumar et al. 2021) utilizing an Agilent Seahorse XFe96 Extracellular Flux Analyzer (Agilent). All cells were optimized for oligomycin concentration (Sigma #O4876), FCCP concentration (Sigma #C2920) and seeding density. Seeding density for the 970, 708 and CUTLL1 cells were 250K, 200K, and 150K cells/well respectively. Final oligomycin concentrations for 970, 708 and CUTLL1 cells were 1μM, 2μM, and 1μM respectively. The final FCCP concentration for 970 and 708 cells was 2μM and CUTTL1 cells were 1uM. Final Rotenone/Antimycin A (Sigma #R8875, #A8674) concentrations for all cell lines were 0.5μM. CellTak (Corning #354240) was used to adhere cells to the plate. Murine T-ALL cells were analyzed at 48 hours after 9nM 4-OHT treatment. Human T-ALL cells were analyzed at 4 days after transduction of shCDC73. Dead cells were excluded by trypan blue staining and protein concentration by Bradford assay (Biorad) was performed to normalize results further to live cell seeding density. Data was analyzed using Wave software (Agilent).

### Metaphase spreads

Murine T-ALL cells were treated with 9nM 4-OHT for 30h. Colcemid (10ug/ml) was added to arrest cells in metaphase. Pre-warmed KCl (37°C) was added by slowly dropping from a Pasteur pipette while gently shaking tube. Cells were fixed in ice cold fixative (3:1 methanol to acetic acid) three times and dropped onto slanted slides from about 10 feet using a Pasteur pipette and humid box. Slides were dried on a slide warmer and visualized under a light microscope to ensure metaphases were visible, then mounted using ProLong Gold Antifade Reagent with DAPI (Life Technologies). Slides were randomized and assigned identification codes independently by a second lab worker and blinded to the first lab worker before imaging on an Olympus BX-61 microscope and viewed with SKYview software (ASI). Blinded images were analyzed by the first lab worker using ImageJ software for chromosomal abnormalities and then unblinded.

### Bru-seq and BruUV-seq library preparation

Cells were split into fresh media upon treatment of the vehicle (ethanol) or OHT. Bru-seq and BruUV-seq were performed as previously described (Paulsen et al. 2013; Paulsen et al. 2014; Magnuson et al. 2015). Briefly, for BruUV-seq, cells were irradiated with 200 J/m^2^ UV-C light. Bromouridine was added at 2mM to label cells in conditioned media directly following UV-irradiation for 30 min. Cells were then lysed directly in TRIzol followed by isolation of total RNA. The isolated total RNA was further treated with DNAse (TURBO DNA-free Kit; Invitrogen) and a spike-in cocktail of Bru-labeled and unlabeled RNA was mixed in with total RNA. Bru-labeled RNA was immunocaptured using anti-BrdU antibodies (BD Biosciences). Anti-BrdU monoclonal antibody was conjugated to magnetic beads and incubated with the RNA. Beads were washed with 200ul 0.1% BSA in PBS and resuspended in 0.1% BSA in PBS and diluted RNaseOUT. Spiked total RNA was added to the conjugated beads and incubated at RT with gentle rotation. Beads were washed, heated at 96°C for 10 min and spun. Beads were captured and supernatant was saved as Bru-RNA. Next, rRNA was removed using FastSelect (Qiagen). First strand cDNA was synthesized via Superscript II, and cleanup was done using AMPure beads. To synthesize second strand, DNA polymerase was added, and cleanup was done using AMPure beads. End Repair was performed with T4 DNA Polymerase, and cleanup was done using AMPure beads. Adenylated 3’ ends were added with Klenow Exo(−)polymerase, and cleanup was done using AMPure beads. Ligation of adaptors was done using the NEB quick ligation kit, and cleanup was done using AMPure beads. Size selection was performed using an agarose gel (3% gel using NuSieve 3:1 agarose). Gel slices were excised at 500bp and purified using the Qiagen QIAEXII kit. Uridine digestion and DNA fragment enrichment was done with USER enzyme. Samples were placed on ice and a universal ligation adapter and dual-index, barcoded primers were added. PCR was performed, and AMPure beads were used to clean up. Pellet was resuspended in 5mM Tris pH 8.0 and incubated at 28 ̊C. Beads were captured, and supernatant was transferred into a low-binding tube.

### Bru-seq and Bru-UV-seq Sequencing and Analysis

Paired-end sequencing of libraries was performed on the NovaSeq 6000 (Illumina) by the University of Michigan Advanced Genomics Core. The resulting fastq reads were trimmed using BBDuk (BB Tools 38.46). Trimmed reads were aligned to the mouse ribosomal RNA (rRNA) repeating unit (GenBank BK000964.3) and mitochondrial genome (from the mm10 reference sequence) using Bowtie2 (2.3.3). Reads that did not align to rRNA or the mitochondrial genome were then aligned to the mouse genome build mm10/GRCm38 using STAR (v 2.5.3a) and a STAR index created from GENCODE annotation version M15 (Frankish et al. 2018, PMID 30357393). Coverages were determined as (1/read length) for each base in the genome using BEDTools2 2.28.0 (Quinlan and Hall 2010). These base coverages were used to obtain pseudo-read counts in strand-specific genomic features, such as genes, using BEDtools intersect. Differential gene expression for Bru-seq was performed using DESeq2 1.18.1 (Love et al. 2014. PMID 25516281). Differentially expressed genes were defined as having fold changes>1.5 (up or down) and adjusted p-values < 0.05. The apeglm method was used for log2 fold change shrinkage (Zhu, et al. 2018). To identify peaks in the BruUV-seq data, plus- and minus-strand read pairs were first separated using Sambamba 0.8.0 and then MACS 2.2.7.1 was used to call peaks and summits for each strand. The MACS2 p-value threshold was set to 0.01 and read duplicates were retained. Differential peak analysis was performed using DiffBind 3.6 (Ross-Innes et al. 2012) and the parameter summits=FALSE was set for the dba.count function to ensure consensus peaks were built around whole peaks from MACS2. These consensus peaks were used for downstream analyses. DESeq2 was the underlying quantification and differential testing method used by DiffBind. DeepTools 3.5.1 was used to generate signal files in bigWig format from uniquely mapped reads. Signal was RPKM-normalized using the total number of uniquely mapped reads in a given library. BruUV-seq peaks were classified in terms of proximity and orientation with respect to TSS and bodies of annotated transcripts, using closestsBed from bedtools 2.30.0, querying TSS coordinates against the 5’ coordinates of BruUV-seq peaks. Only protein-coding transcripts were used, and genes were required to be expressed (mean Bru-seq RPKM>0.25 across samples). Where TSS coordinates are identical, the longer transcript was used. Peaks were classified as “TSS” if within 2 kb of the closest TSS, “gene body” if overlapping gene (sense orientation only), or “intergenic” if not classified as TSS or gene body (this includes peaks antisense to expressed genes). To create the eRNA heatmaps (Fig. S7D-E), normalized counts data and peak data were filtered for intergenic peaks and antisense gene body peaks. Data was further subset into downregulated genes using cutoff of FDR<0.05 and FC<0; upregulated genes had cutoff of FDR<0.05 and FC>0. These subsets of downregulated and upregulated genes were log transformed, then used to generate a heatmaps separated by cell line using R package pheatmap (1.0.12).

### ChIP-PCR, library preparation and sequencing

ChIP-PCR and ChIP-Seq library preparation and sequencing was performed as previously described (McCarter et al. 2020) with the following additions. All reactions were followed by a SPRI bead cleanup using AMPure or SPRISelect beads. Size selection for the library was performed by using AMPure or SPRISelect selection. Paired-end sequencing of H3K27ac, H2BKUb1, CDC73, and ETS1 ChIP libraries and unenriched control input libraries were performed using Nova-seq or Next-seq with approximately 20M reads per sample.

### ChIP-Seq analysis

ChIP-seq alignment, filtering, track generation, peak calling, overlaps, and differential binding analyses were performed as previously published (McCarter et al. 2020). The exception was that for mouse cells the reads were aligned to the mm10 genome assembly and post filtered for known ENCODE blacklist regions in mice. For differential H3K27ac peak analysis in OHT vs. vehicle treated 969, 970, and 708 cells, DiffBind 3.6 was used (parameter summits=FALSE). DESeq2 was chosen for the analysis method and adjusted p-value< 0.05 was used as the significance threshold. Consensus peaks from DiffBind were used in downstream analyses. Signal files in bigWig format were produced with bamCoverage (DeepTools 3.5.1). Intergenic peaks (>2 kb from expressed gene TSS) were centered and the average (unstranded) signal was plotted for given samples +/−2.5 kb from the center using computeMatrix and plotProfile (DeepTools; Fig 6E, Fig S5C). For differential H2BK120ub1 peak analysis comparing control versus OHT, HOMER analyzeRepeats.pl was first run on mm10 genes to count tags on the full gene body (TSS to TTS) in a strand-insensitive manner, with parameters: -raw -gid -count genes -strand both. Results were then passed to HOMER getDiffExpression.pl which used DESeq2 to quantify and test differential histone ubiquitination across OHT vs EtOH comparisons in each cell line. Metagene plots were created using DeepTools for all genes or OXPHOS and DNA repair genes (Table S1). The bigwig display files were used as the input file scores to be plotted. The computeMatrix (“scale-regions”) tool was used to generate a score per genome matrix −1kb from the TSS and +1kb from the TTS. This matrix was then input into plotProfile to create the metagene plot of scores over genes. H3K27Ac and BruUV-seq consensus peaks were integrated using closestBed (bedtools), querying H3K27Ac peaks against BruUV-seq 5’ coordinates and requiring peaks to overlap (distance = 0). Dynamic intergenic H3K27ac/eRNA peak pairs were defined as having adjusted p-values<0.05 (OHT/control) in both assays and in both 969 and 970 cells. As per BruUV-seq peak classification described above, the “intergenic” definition included peaks that were >2 kb from expressed gene TSS and those antisense to gene bodies. CDC73 signal was quantified over gene bodies using multiBamCov (bedtools), where genes were defined by the human GENCODE version 27 basic annotation and uniquely mapped reads (mapping quality>30). RPKM was computed using the total number of uniquely mapped reads in the original alignments (mapping quality>30). ETS1 and RBPJ promoter regions were quantified in the same manner as CDC73, except limited to the first 2kb of each gene.

## Additional statistical information

Unless otherwise indicated, P-values were derived from two-sided two-sample t-tests of Log2-transformed data for comparisons in experiments involving two groups and 1-way ANOVA of Log2-transformed data for pairwise contrasts in experiments with more than two groups. Unless otherwise stated, horizontal lines are means and values are shown as mean ± standard deviation. Survival curves (or time to event data) was tested with log-rank tests comparing pairs of groups. Gene set enrichment analysis (GSEA) was performed using GSEAv4.1.0 (Broad Institute) and gene lists from MSigDBv7.4 and custom lists of high confidence human direct NOTCH1 and ETS1 target gene signatures in T-ALL (Wang et al. 2014; McCarter et al. 2020).

## Acknowledgements

We thank Qing Li, David Ferguson, Shannon Carty, Michelle Paulsen, Arvind Rao, Rohan Kogdule, Amparo Serna Alarcon, Ran Yan, Cher Sha, Morgan Jones, Koral Campbell, Andrea Hartlerod, Courtney Hames, and Luis Correa for their thoughtful input and technical assistance during this project. We thank Andrew Weng, Malathi Kandarpa, and Moshe Talpaz for PDX and primary T-ALL samples. We thank Daniela Salgado Figueroa and Katia Georgopoulos for Hi-C contact maps of GSE79422 (Kashiwagi et al. 2022). This work was supported by funding from the National Institutes of Health and National Cancer Institute (NIH/NCI 1F31CA260929 [AM], NIH/NCI R01CA196604 [MC], NIH/NIAID R01 AI136941-01 [MC], NIH T-32-GM007315 [AM], and R01GM101171 [DL]), The University of Michigan Rackham Graduate School, Michigan Medicine Rogel Cancer Center, Rally Foundation for Childhood Cancer Research, and Alex’s Lemonade Stand Foundation.

## Author Contributions

Conceptualization: M.Y.C., A.F.M. Investigation: A.F.M, S.L., N.A.D., A.W., E.C., B.M., A.C.M, F.A., N.K, C.B., G.B., Y.L., Q.W. Visualization: M.Y.C., A.F.M, S.L., N.D., E.C., S.K., D.B.L., M.L., A.C.M., J.S., K.L., R.J.H.R. A.G.M. Formal Analysis: A.F.M., E.C., B.M., N.A.D. Data Curation: A.C.M, N.K., K.L., E.C., B.M., M.Y.C., R.J.H.R. Funding acquisition: M.Y.C., A.F.M. Writing of the original draft: A.F.M., M.Y.C. Review and editing of the manuscript: M.Y.C, A.F.M. R.J.H.R., D.B.L., M.L., S.K., B.M., E.C., N.K., K.L., N.A.D., J.S., C.M., A.G.M. Supervision: M.Y.C.

