## Supplemental figures, tables, and methods for "Cdc73 protects Notch-induced T-cell leukemia cells from DNA damage and mitochondrial stress"

**List of Supplemental Materials**

Figure S1 -- CDC73 interacts with NOTCH1 and ETS1 in T-ALL cells

Figure S2 -- *CDC73* is expressed throughout T-cell development

Figure S3 -- *CDC73* is expressed in diverse oncogenomic T-ALL subsets

Figure S4 -- Cdc73 shares Notch- and ETS1-driven pathways

Figure S5 -- Cdc73 is important for genome integrity

Figure S6 -- Cdc73 is important for oxidative phosphorylation

Figure S7 -- Cdc73 does not primarily promote DNA repair and OXPHOS gene expression through enhancers

Figure S8 -- Cdc73 promotes DNA repair and OXPHOS gene expression through canonical mRNA functions at gene bodies

Figure S9 -- Cdc73, Notch, and Ets1 signals intersect at gene expression to mitigate metabolic and genotoxic stresses of elevated Notch signals

Table S1 – Lists of DNA repair and OXPHOS core enrichment genes found in GSEA analysis of $Cdc73^{f/f}$ T-ALL cell lines

Table S2 – Lists of antibodies and primers

Supplemental Material and Methods

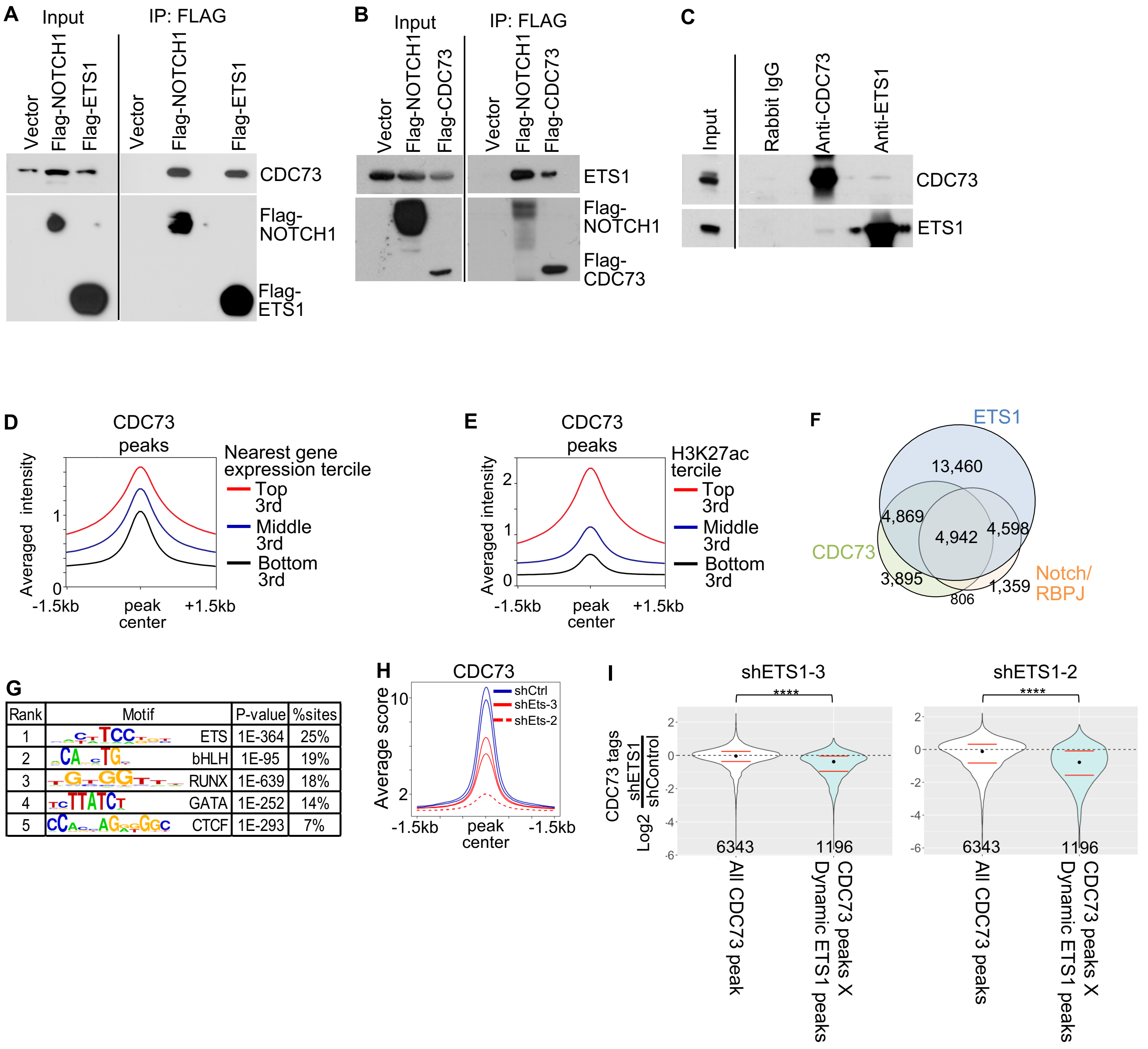

**Figure S1. CDC73 interacts with NOTCH1 and ETS1 in T-ALL cells.** A) Co-IP with Flag-NOTCH1 and Flag-ETS1 showing interactions between NOTCH1 or ETS1 with CDC73 in CEM T-ALL cells treated with benzonase. B) Co-IP with Flag-CDC73 showing interaction of ETS1 with CDC73 in 8946 T-ALL cells treated with benzonase. C) Endogenous co-IP showing interaction between ETS1 and CDC73 in THP-6 T-ALL cells treated with benzonase. D-E) Metagene plots of averaged CDC73 ChIP-Seq tags of two THP-6 bioreplicates at ATAC-Seq peaks ranked by tercile mean FPKM expression of nearest genes (D; N=4) or tercile mean H3K27ac tags (E; N=2) (McCarter et al. 2020). F) Venn diagram showing overlap of ETS1, CDC73, and Notch/RBPJ ChIP-Seq peaks in THP-6 cells. G) De novo motif analysis of merged CDC73 ChIP-Seq peaks in THP-6 cells from (McCarter et al. 2020). H) Metagene plot of averaged CDC73 ChIP-Seq peaks that overlap with dynamic ETS1 peaks in human THP-6 cells transduced with either control shRNA (solid blue line; N=2) or shETS1-3 (solid red line; N=2) or shETS1-2 (dotted red line; N=2). Dynamic ETS1 peaks were defined as ETS1 peaks that diminish with FDR<0.1 upon knockdown with two independent shETS1 (McCarter et al. 2020). I) Violin plots showing the CDC73 ChIP-Seq Log2FC for all CDC73 peaks and CDC73 peaks that overlap with dynamic ETS1 peaks in context of ETS1 knockdown with shETS1-3 (left) or shETS1-2 (right).

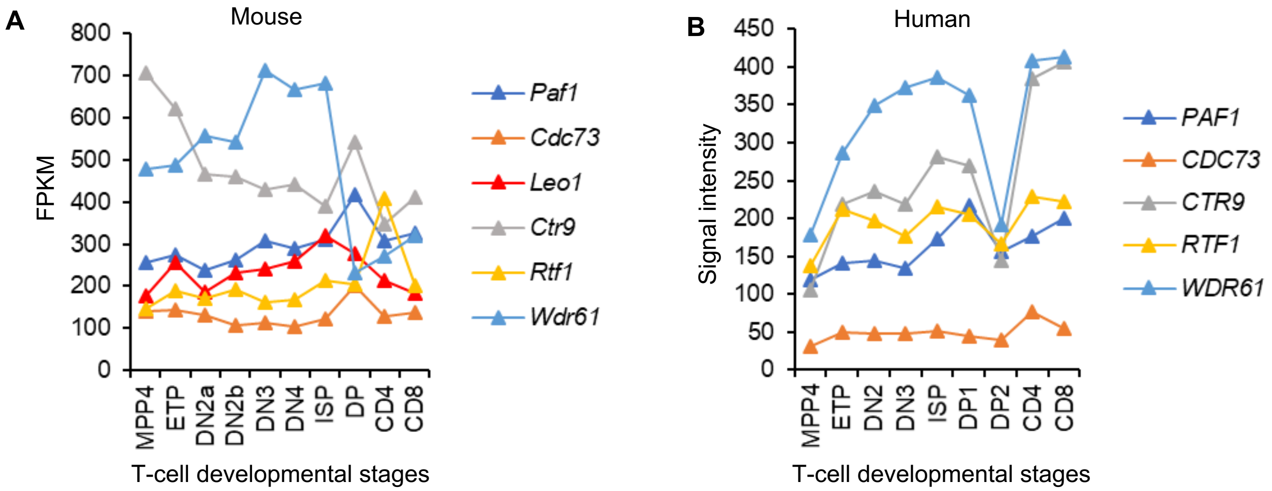

**Figure S2. *CDC73* is expressed throughout T-cell development.** A) Expression of Paf1C family members in murine T-cell developmental subsets (ImmGen; GSE109125). B) Expression of PAF1C family members in human T-cell development (GSE22601).

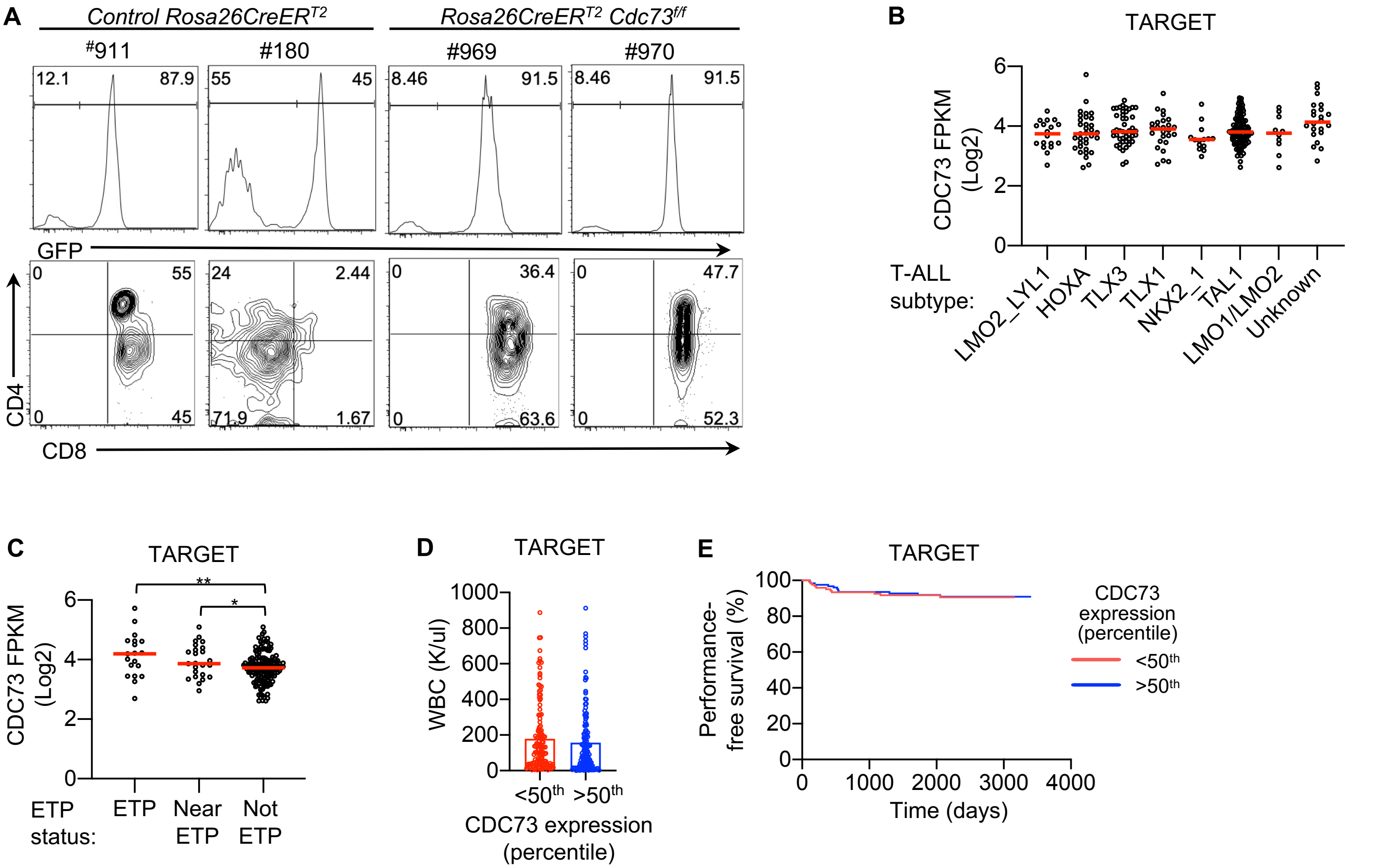

**Figure. S3. *CDC73* is expressed in diverse T-ALL subsets.** A) Representative CD4/CD8 flow cytometric profiles of GFP+ splenic tumors induced by activated Notch1 alleles (*ΔE/Notch1*) in *Rosa26CreER$^{T2}$* control and *Rosa26CreER$^{T2}$ Cdc73$^{f/f}$* mice. B-C) *CDC73* expression according to T-ALL subtype (B) and ETP status (C) in the TARGET database. D) White blood cell (WBC) counts of T-ALL patients stratified by CDC73 expression. E) Survival curves of T-ALL patients stratified by *CDC73* expression. TARGET=Therapeutically Applicable Research to Generate Effective Treatments (TARGET) (https://ocg.cancer.gov/programs/target) initiative, phs000218 (Liu et al. 2017). The ALL project team was headed by Stephen P. Hunger, M.D. at the University of Colorado Cancer Center, Denver, CO, USA. The dbGaP Sub-study ID was phs000463/phs000464. The data used for this analysis are available at https://portal.gdc.cancer.gov/projects.

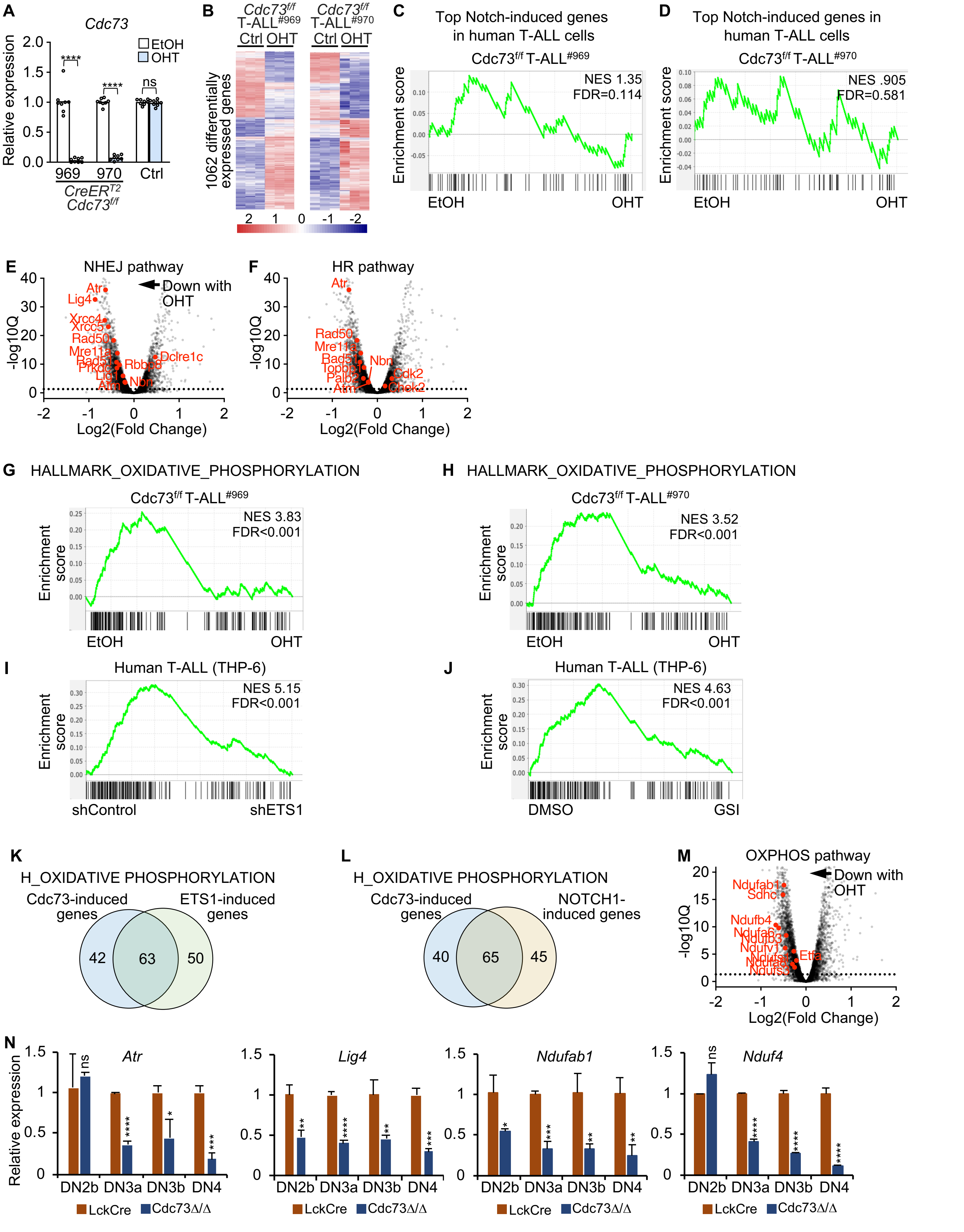
A
Cdc73
Relative expression
EtOH
OHT
****
****
ns
969
970
Ctrl
CreERT2
Cdc73f/f
B
Cdc73f/f T-ALL#969
Cdc73f/f T-ALL#970
Ctrl OHT
Ctrl OHT
1062 differentially expressed genes
C
Top Notch-induced genes in human T-ALL cells
Cdc73f/f T-ALL#969
Enrichment score
NES 1.35
FDR=0.114
EtOH
OHT
D
Top Notch-induced genes in human T-ALL cells
Cdc73f/f T-ALL#970
NES .905
FDR=0.581
E
NHEJ pathway
Down with OHT
-log10Q
Log2(Fold Change)
F
HR pathway
G
HALLMARK_OXIDATIVE_PHOSPHORYLATION
Cdc73f/f T-ALL#969
NES 3.83
FDR<0.001
H
HALLMARK_OXIDATIVE_PHOSPHORYLATION
Cdc73f/f T-ALL#970
NES 3.52
FDR<0.001
I
Human T-ALL (THP-6)
NES 5.15
FDR<0.001
shControl
shETS1
J
Human T-ALL (THP-6)
NES 4.63
FDR<0.001
DMSO
GSI
K
H_OXIDATIVE PHOSPHORYLATION
Cdc73-induced genes
ETS1-induced genes
42
63
50
L
H_OXIDATIVE PHOSPHORYLATION
Cdc73-induced genes
NOTCH1-induced genes
40
65
45
M
OXPHOS pathway
Down with OHT
N
Atr
Lig4
Ndufab1
Nduf4
Relative expression
DN2b DN3a DN3b DN4
LckCre
Cdc73Δ/Δ

**Figure S4. Cdc73 shares Notch- and ETS1-driven pathways**. A) Relative expression of *Cdc73* in the 969 and 970 *Rosa26CreER*$^{T2}$ *Cdc73*$^{f/f}$ ∆E/Notch1-induced T-ALL cell lines and control *Rosa26CreER*$^{T2}$ ∆E/Notch1-induced T-ALL cell line after treatment with OHT for 30 hours. B) Heatmap of 1062 differentially expressed genes upon *Cdc73* deletion shared between the 969 and 970 cell lines. C-D) GSEA using the direct Notch target signature in T-ALL (Wang et al., 2014) of Cdc73-induced genes. E-F) Volcano plots of significance vs. Bru-Seq Log2FC data showing the control vs. OHT (Cdc73∆/∆) comparison and important genes in the non-homologous end-joining (NHEJ; E) and homologous recombination (HR; F) pathways in 970 T-ALL cells on background of all genes (grey) giving average RPKM>0.8. G-J) GSEA using the MSigDB Hallmark_Oxidative_Phosphorylation gene list of Cdc73-induced genes in 969 cells (G), Cdc73-induced genes in 970 cells (H), ETS1-induced genes in THP-6 cells (McCarter et al., 2020) (I), and Notch-induced genes in THP-6 cells (McCarter et al., 2020) (J). K-L) Venn diagram showing overlap of Cdc73-induced and ETS1-induced (K) or Notch-induced (L) genes in the Hallmark_Oxidative_Phosphorylation gene list. M) Volcano plot of significance vs. Bru-Seq Log2FC data showing the control vs. OHT (Cdc73∆/∆) comparison and important genes in the oxidative phosphorylation pathways in 970 T-ALL cells on background of all genes (grey) giving average RPKM>0.8. N) Relative expression of important DNA repair (*Atr* and *Lig4*) and oxidative phosphorylation (*Ndufab1* and *Nduf4*) genes in sorted thymic subsets from *LckCre* mice (red) and *Cdc73*$^{\Delta/\Delta}$ mice (blue).

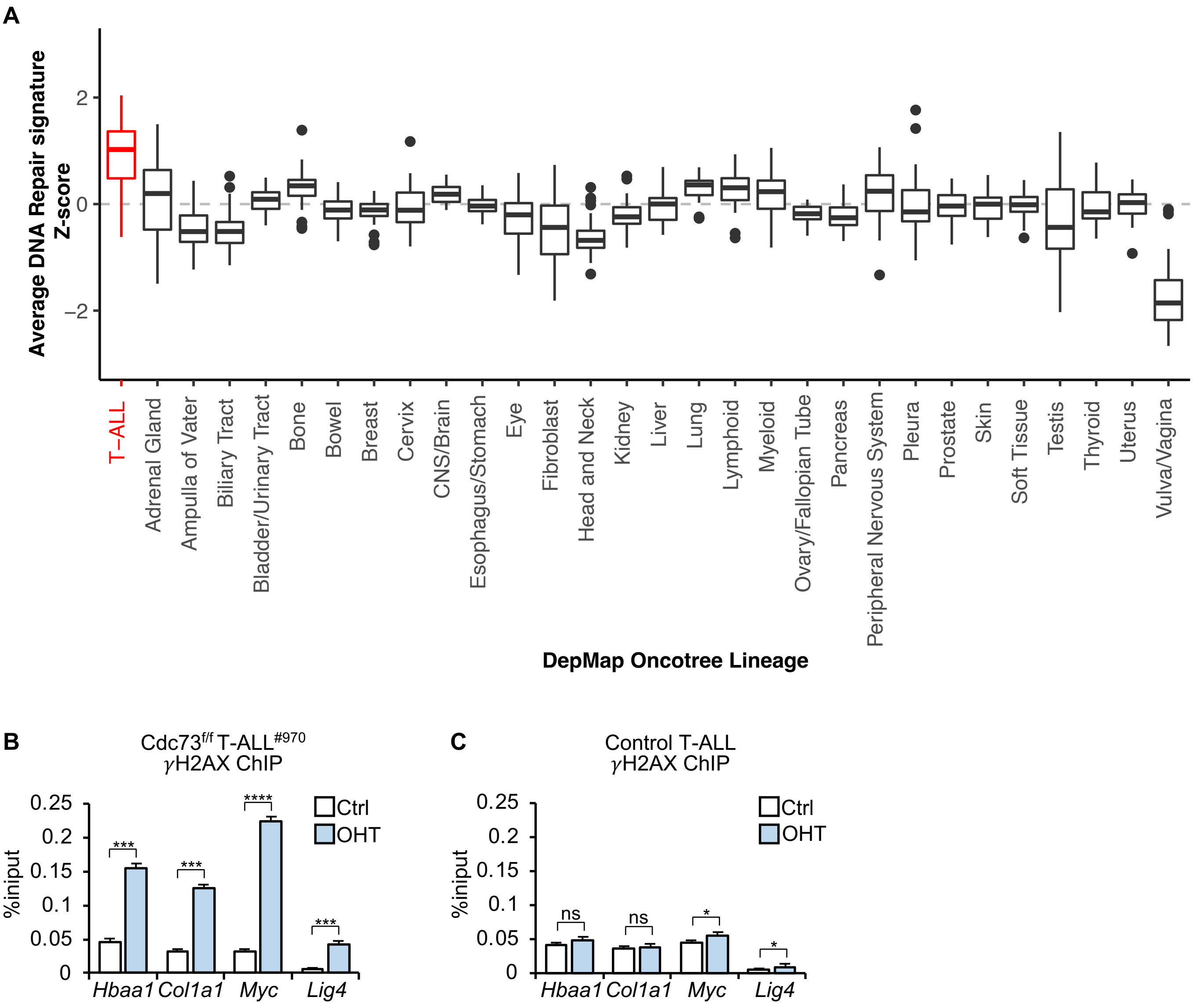

**Figure S5. Cdc73 is important for genome integrity**. A) Average z-scores of RNA-Seq expression of 34 DNA repair target genes induced by Cdc73, ETS1, and Notch based on GSEA core enrichment analysis using the Kauffman_DNA_repair gene list (Table S1) across all cell lines grouped by Oncotree lineage in DepMap 22Q4. B-C) γH2AX qChIP in *$Cdc73^{f/f}$* T-ALL cells (970, B) and control *$Rosa26CreER^{T2}$* T-ALL cells (C) treated with OHT for 30 hours.

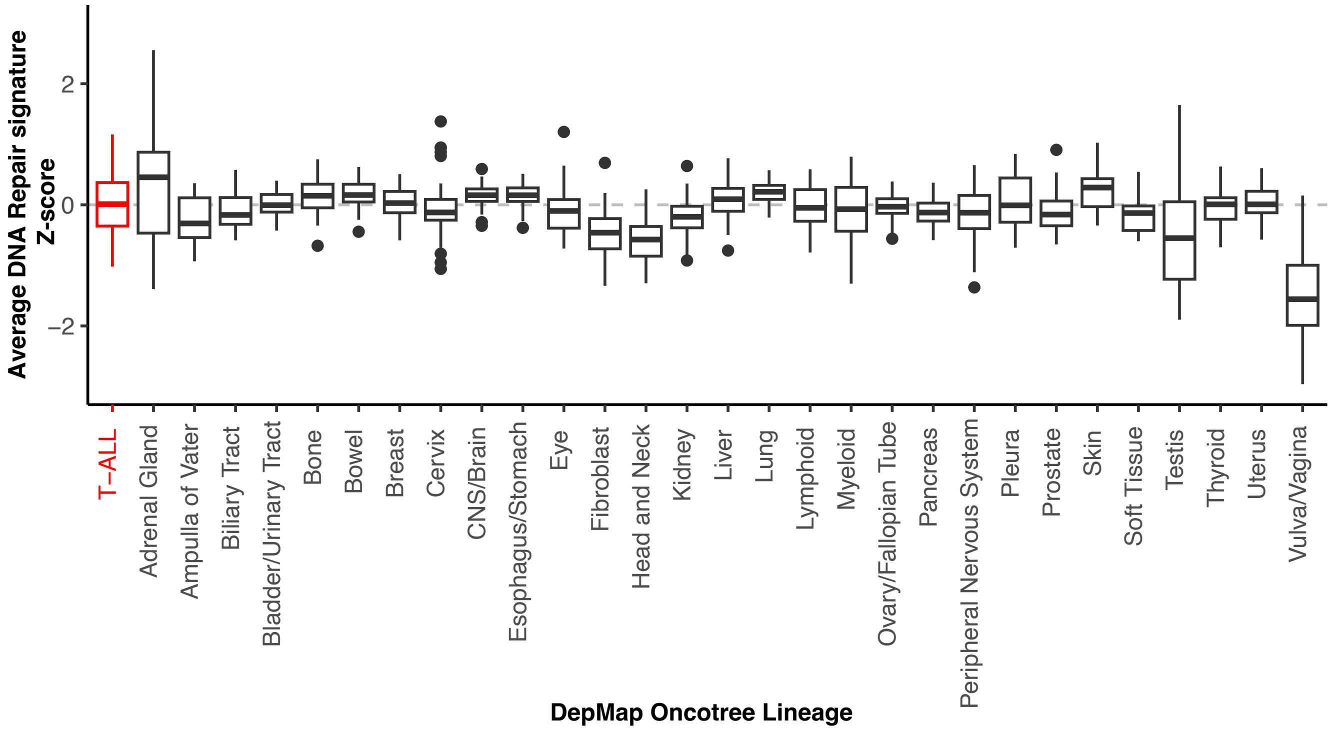

**Figure S6. Cdc73 is important for oxidative phosphorylation.** Average z-scores of RNA-Seq expression of 38 OXPHOS genes induced by Cdc73, ETS1, and Notch based on GSEA core enrichment analysis using the Hallmark_oxidative_phosphorylation gene list across all cell lines grouped by lineage in DepMap 22Q4.

Melnick_Fig.S7

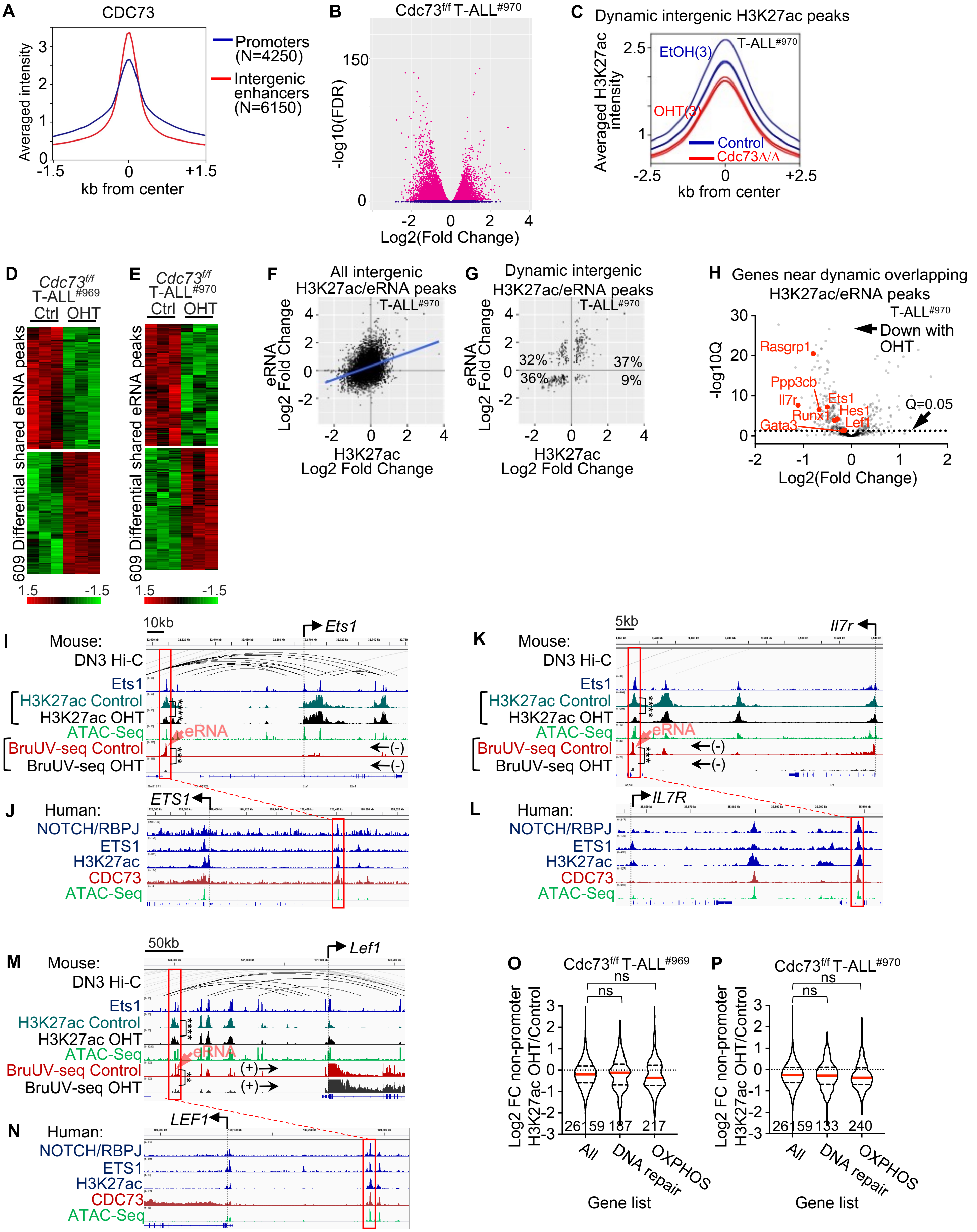

**Figure S7. Cdc73 does not primarily promote DNA repair and OXPHOS gene expression through enhancers.** A) Metagene plots of intergenic and promoter CDC73 ChIP-Seq signals in THP-6 cells. B) Volcano plot of significance vs. Log2(Fold Change OHT/Control) H3K27ac ChIP-Seq signals of *Rosa26CreER$^{T2}$ Cdc73$^{f/f}$* T-ALL cells (970) upon treatment with 6nM OHT for 30 hours. C) Metagene plot of dynamic intergenic H3K27ac signals (FDR<0.05 in 969 and 970 cells but not in control cells) in 970 cells. D-E) Heatmaps of differential eRNA peaks shared between the 969 (E) and 970 (E) cell lines. eRNAs were defined as intergenic BruUV-Seq peaks or intragenic peaks that were antisense in direction relative to mRNAs. F-G) BruUV-Seq Log2(OHT/Control) versus H3K27ac Log2(OHT/Control) scatterplots of all overlapping intergenic peaks (F) or overlapping dynamic intergenic peaks (G) in *Cdc73$^{f/f}$* 970 T-ALL cells. Overlapping "dynamic peaks" were defined as giving q<0.05 and FDR<0.05 in the same direction for the BruUV-Seq and H3K27ac comparisons respectively in both *Cdc73$^{f/f}$* T-ALL cells but not in control T-ALL cells. H) Volcano plot of significance vs. Bru-Seq Log2(OHT/Control) of genes nearest OHT-downregulated dynamic intergenic BruUV-Seq and H3K27ac overlapping peaks in 970 *Cdc73$^{f/f}$* T-ALL cells. I-N) Display tracks of indicated ChIP-Seq and ATAC-seq datasets at the *Ets1* (I-J), *Il7r* (K-L), and *Lef1* (M-N) loci in mouse 969 cells (I, K, M) or human THP-6 cells (J, L, N) showing nearest mouse-human homologous enhancers in red boxes that contain overlapping dynamic intergenic eRNA and H3K27ac peaks. Ets1 ChIP-seq (GSM461516); ATAC-seq (GSM2461649); DN3 Hi-C (GSE79422) analyzed in (Kashiwagi et al. 2022). O-P) Violin plots showing H2K27ac ChIP-Seq Log2FC in 969 cells (O) and 970 cells (P) at non-promoter H2K27ac peaks nearest all genes, DNA repair genes, and OXPHOS genes in the GSEA enrichment cores of 969 and 970 cells (Table S1). OHT was added for 30 hours to delete *Cdc73.* Numbers below violin plots represent number of genes. **FDR<0.01; ***FDR<0.001; ****FDR<0.0001.

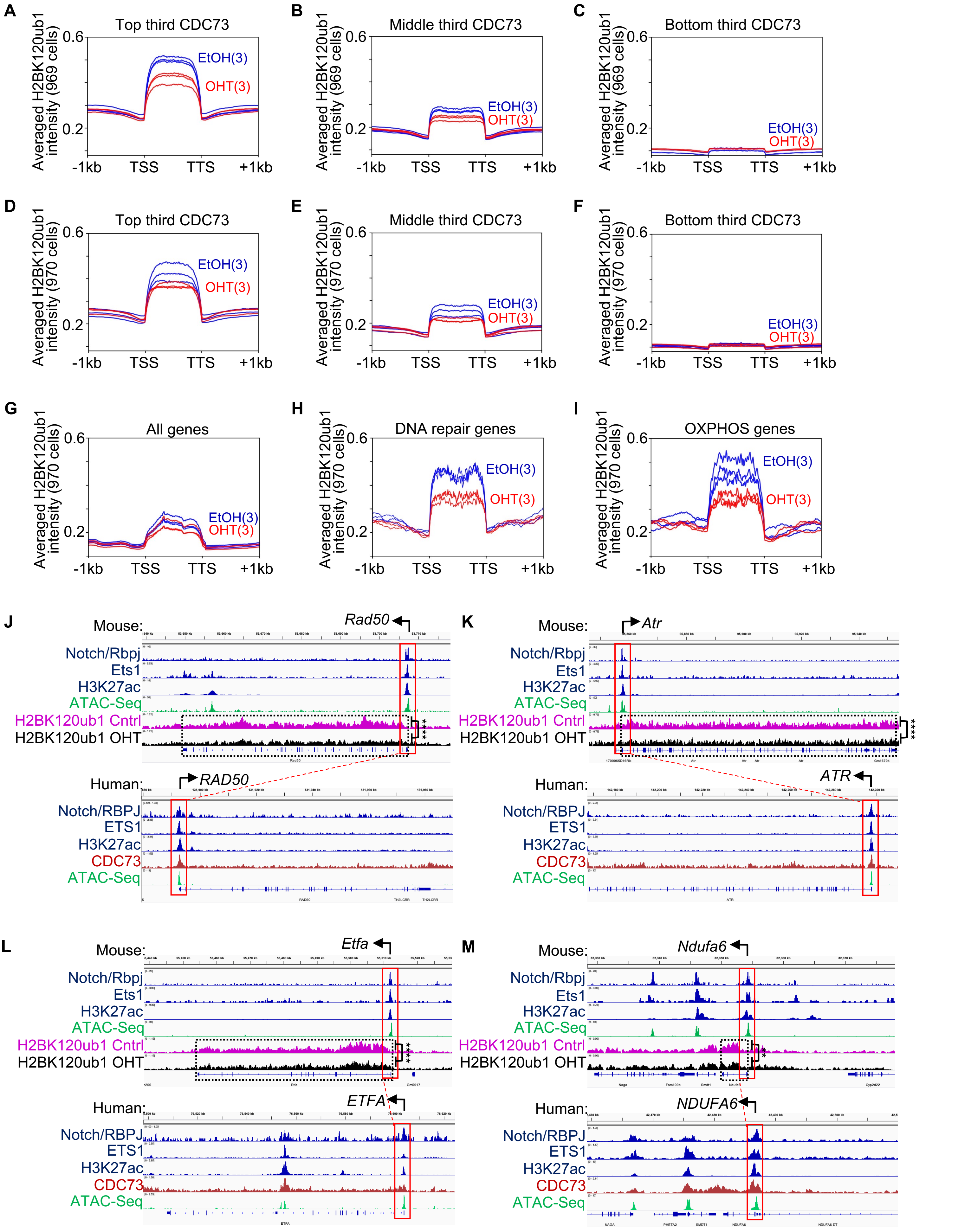
A
Top third CDC73
Averaged H2BK120ub1 intensity (969 cells)
EtOH(3)
OHT(3)
B
Middle third CDC73
C
Bottom third CDC73
D
E
F
Averaged H2BK120ub1 intensity (970 cells)
G
All genes
H
DNA repair genes
I
OXPHOS genes
0.6
0.2
-1kb
TSS
TTS
+1kb
J
Mouse:
Rad50
Notch/Rbpj
Ets1
H3K27ac
ATAC-Seq
H2BK120ub1 Cntrl
H2BK120ub1 OHT
Human:
RAD50
Notch/RBPJ
ETS1
H3K27ac
CDC73
ATAC-Seq
K
Atr
ATR
L
Etfa
ETFA
M
Ndufa6
NDUFA6

**Figure S8. Cdc73 promotes DNA repair and OXPHOS gene expression through canonical mRNA functions at gene bodies.** A-F) Metagene plots of H2BK120ub1 signals in EtOH (blue) and OHT (red) treated 969 cells (A-C) and 970 cells (D-F) at genes with CDC73 signals of human homologs in THP-6 cells ranked at the top tercile (A,D), middle tercile (B,E), and bottom tercile (C,F). G-I) Metagene plots of H2BK120ub1 signals in EtOH- (blue) and OHT- (red) treated 970 cells at all genes (G), DNA repair genes (H), and OXPHOS genes (I) in core enrichment genes of GSEA analyses (Fig. 3J, Fig. S4H, Table S1). J-M) Display tracks of indicated ChIP-Seq and ATAC-Seq datasets at DNA repair genes (J-K) and OXPHOS genes (L-M) in mouse 969 cells (top) or human THP-6 cells (bottom) showing representative tracks and FDR values upon OHT addition (*Cdc73* deletion) of H2K120ub1 signals between TSS and TTS. **FDR<0.01. ****FDR<0.0001. ATAC-seq (GSM2461649). Ets1 ChIP-Seq (GSM2461515).

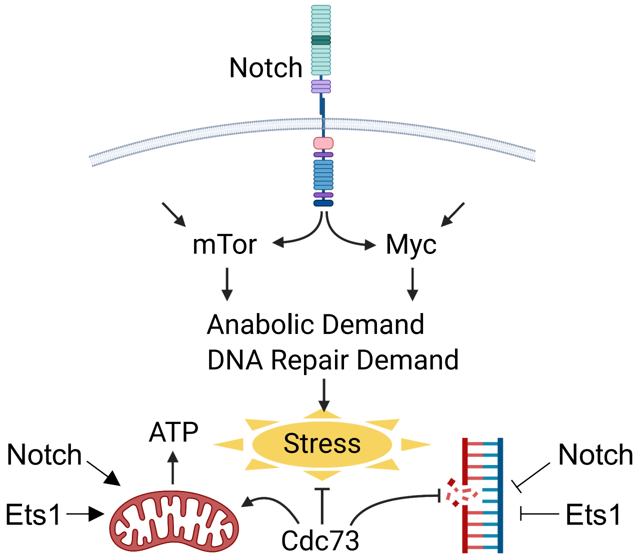

**Figure S9. Cdc73, Notch, and Ets1 signals intersect at gene expression to mitigate metabolic and genotoxic stresses of elevated Notch signals.** Elevated Notch signals induce major oncogenic pathways, mTorc1 and Myc, which increases demand for cellular energy and DNA repair. Notch, Ets1, and Cdc73 induce mRNA synthesis of a subset of highly expressed genes in DNA repair and oxidative phosphorylation pathways to support elevated and oncogenic Notch functions. Generated using Biorender.

### Supplemental Material and Methods

#### Differential analysis of CDC73 signals comparing control and ETS1 knockdown conditions

Active nucleosome free regions (NFRs) in THP-6 cells were defined as ATAC-Seq peaks associated with the top one-third of H3K27ac signal within a +/-4kb window (15,000 intervals) based on previously published datasets (McCarter et al. 2020). Intersect function from BEDtools 2.30.0 (Quinlan and Hall 2010) (https://github.com/arq5x/bedtools2/blob/master/README.md) was used to intersect active NFR intervals with control CDC73 ChIP-seq peaks from two experiments. Intersected bed files were concatenated to find all intervals where control CDC73 peaks overlap with active NFR intervals. We performed differential binding analysis using Diffbind 3.6 (Ross-Innes et al. 2012) (https://bioconductor.org/packages/release/bioc/html/DiffBind.html) and EdgeR algorithm (Robinson et al. 2010; McCarthy et al. 2012) (https://bioconductor.org/packages/release/bioc/html/edgeR.html) to compare CDC73 binding in control (N=2) vs. shETS1-2 (N=2), and shETS1-3 (N=2) samples. We designated the previously identified intervals where control CDC73 peaks overlap with active NFRs as "all CDC73 peaks". We defined "dynamic ETS1 peaks" as intervals where ETS1 binding decreases (FC<0) with FDR<0.1 for both shETS1-2 and shETS1-3 (McCarter et al. 2020). We intersected these intervals with "all CDC73 peaks" and designated resulting intervals as "CDC73 peaks X dynamic Ets1 peaks." Violin plots were generated to compare CDC73 binding fold change at "all CDC73 intervals" and "CDC73 peaks X dynamic Ets1" for shETS1-2 and shETS1-3 treated samples. Plots also display median, 1$^{st}$ and 3$^{rd}$ quartiles. Using deepTools Galaxy computeMatrix and plotHeatmap(Ramirez et al. 2016), we generated metagene plots at "CDC73 x dynamic Ets1 peaks" intervals using ChIP-Seq reads from CDC73 shControl, CDC73 shETS1-2, and CDC73 shETS1-3 bigwig files (McCarter et al. 2020).

**Generation of z-scores for DNA repair and OXPHOS gene lists from DepMap cell line RNA-Seq datasets.** (The cell line subclassification ("Model.csv") and RNA-Seq ("OmicsExpressionProteinCodingGenesTPMLogp1.csv") data matrices from DepMap Public 22Q4 files were read into R (v4.2.2). These matrices were subsetted and merged using dplyr (v1.0.10) to create a final main data frame containing ACH-ID, stripped cell line name, Oncotree lineage, and gene expression values that are found in core enrichment gene lists in Kauffman_DNA_repair and Hallmark_Oxidative_Phosphorylation shared between Cdc73-induced target genes in 969 and 970 cells, Notch-induced target genes in THP-6 cells, and ETS1-induced target genes in THP-6 cells (Table S2). This dataframe was scaled using scale() to standardize genes to z-score. The columns containing the scaled gene expression values were then averaged to obtain a gene score for the gene sets in each cell line. The T-ALL group was subset based on stripped cell line name, while the other groups of interest for comparison were subset based on Oncotree lineage. These values were plotted using ggplot2 (v3.3.6).

Table S1. Core enrichment genes in Kaufmann_DNA_repair for Cdc73-induced target genes ir

| RANK | SYMBOL |
|---|---|
| 1 | LIG4 |
| 2 | RAD51B |
| 3 | XRCC4 |
| 4 | FANCD2 |
| 5 | FANCL |
| 6 | NEIL3 |
| 7 | CETN3 |
| 8 | ATR |
| 9 | WRN |
| 10 | RAD18 |
| 11 | XRCC5 |
| 12 | RBBP8 |
| 13 | SMC2 |
| 14 | PRKDC |
| 15 | RAD54B |
| 16 | POLE |
| 17 | PMS1 |
| 18 | RPA3 |
| 19 | MBD4 |
| 20 | RAD51AP1 |
| 21 | TOP3A |
| 22 | PARP2 |
| 23 | RAD51 |
| 24 | BRIP1 |
| 25 | MSH2 |
| 26 | CCNH |
| 27 | RFC4 |
| 28 | FANCB |
| 29 | MRE11 |
| 30 | DCLRE1B |
| 31 | MLH1 |
| 32 | EME1 |
| 33 | TDP1 |
| 34 | POLA1 |
| 35 | N4BP2 |
| 36 | POLK |
| 37 | TOPBP1 |
| 38 | FANCC |
| 39 | DDB2 |
| 40 | RAD54L |
| 41 | SMC1A |
| 42 | NUDT1 |
| 43 | TOP2B |

| | |
|---|---|
| 44 | TDG |
| 45 | BLM |
| 46 | ATM |
| 47 | MLH3 |
| 48 | ALKBH3 |
| 49 | TEP1 |
| 50 | RFC1 |
| 51 | RAD1 |
| 52 | XPC |
| 53 | POLQ |
| 54 | CHEK1 |
| 55 | POLD1 |
| 56 | GTF2H4 |
| 57 | MTOR |
| 58 | MBD5 |
| 59 | NBN |
| 60 | TERF1 |
| 61 | RAD51C |
| 62 | UBE2V2 |
| 63 | ERCC6 |
| 64 | RAD51D |
| 65 | XPA |
| 66 | RAD52 |
| 67 | FANCE |
| 68 | UBE2I |
| 69 | DUT |
| 70 | RAD17 |
| 71 | HMGB2 |
| 72 | ATRX |
| 73 | CRY1 |
| 74 | ENDOV |
| 75 | APEX2 |
| 76 | MSH3 |
| 77 | TOP3B |
| 78 | RECQL5 |
| 79 | CDK7 |
| 80 | MPG |
| 81 | RFC2 |
| 82 | FEN1 |
| 83 | MBD2 |
| 84 | RFC3 |
| 85 | FANCA |
| 86 | LIG1 |
| 87 | WDR33 |
| 88 | CRY2 |

| | |
|---|---|
| 89 | EXO1 |
| 90 | APTX |
| 91 | MSH6 |
| 92 | RAD23A |
| 93 | SMC3 |
| 94 | TP53 |
| 95 | UBE2D3 |
| 96 | BRCA2 |
| 97 | ERCC4 |
| 98 | RFC5 |
| 99 | SMC4 |
| 100 | RAD21 |
| 101 | POLE4 |
| 102 | SSRP1 |
| 103 | XRCC3 |
| 104 | RRM1 |
| 105 | MBD1 |
| 106 | ALKBH2 |
| 107 | GTF2H3 |
| 108 | REV3L |
| 109 | MGMT |
| 110 | SUPT16H |
| 111 | PARP4 |
| 112 | CETN2 |

Table S1. Core enrichment genes in Kaufmann_DNA_repair for Cdc73-induced target genes i

| RANK | SYMBOL |
|---|---|
| 1 | LIG4 |
| 2 | FANCL |
| 3 | CETN3 |
| 4 | FANCD2 |
| 5 | XRCC4 |
| 6 | RAD51B |
| 7 | ATR |
| 8 | WRN |
| 9 | XRCC3 |
| 10 | POLD1 |
| 11 | XRCC5 |
| 12 | RAD18 |
| 13 | NEIL3 |
| 14 | NUDT1 |
| 15 | TOP3A |
| 16 | EME1 |
| 17 | RAD54L |
| 18 | POLE |
| 19 | RPA3 |
| 20 | FANCB |
| 21 | DCLRE1B |
| 22 | RFC4 |
| 23 | MBD4 |
| 24 | FANCE |
| 25 | MLH1 |
| 26 | PARP2 |
| 27 | MSH2 |
| 28 | ENDOV |
| 29 | PMS1 |
| 30 | RAD54B |
| 31 | CCNH |
| 32 | RAD51AP1 |
| 33 | RAD23A |
| 34 | RUVBL2 |
| 35 | PRKDC |
| 36 | RAD51 |
| 37 | RFC2 |
| 38 | MRE11 |
| 39 | UBE2I |
| 40 | DUT |
| 41 | MTOR |
| 42 | RFC5 |
| 43 | DDB2 |

| | |
|---|---|
| 44 | TOP3B |
| 45 | ALKBH3 |
| 46 | RAD52 |
| 47 | XPC |
| 48 | TDP1 |
| 49 | FANCC |
| 50 | SMC2 |
| 51 | RAD1 |
| 52 | RBBP8 |
| 53 | TOPBP1 |
| 54 | FANCA |
| 55 | GTF2H4 |
| 56 | MUS81 |
| 57 | TDG |
| 58 | ALKBH2 |
| 59 | NEIL1 |
| 60 | ERCC4 |
| 61 | GTF2H3 |
| 62 | CRY2 |
| 63 | POLD2 |
| 64 | MMS19 |
| 65 | LIG1 |
| 66 | XPA |
| 67 | MBD3 |
| 68 | ERCC6 |
| 69 | APTX |
| 70 | BLM |
| 71 | UBA1 |
| 72 | BRIP1 |
| 73 | LIG3 |
| 74 | NBN |
| 75 | CETN2 |
| 76 | TOP2B |
| 77 | ATM |
| 78 | FEN1 |
| 79 | TERT |
| 80 | POLK |
| 81 | RECQL5 |
| 82 | FANCG |
| 83 | POLA1 |
| 84 | N4BP2 |
| 85 | SMC1A |
| 86 | CDK7 |
| 87 | POLM |
| 88 | RAD9A |

| | |
|---|---|
| 89 | TP53 |
| 90 | RAD51C |
| 91 | MLH3 |
| 92 | MPG |
| 93 | EXO1 |
| 94 | RAD17 |
| 95 | CHAF1A |
| 96 | NEIL2 |
| 97 | RAD51D |

Table S1. Core enrichment genes in Kaufmann_DNA_repair list that are shared between Cdc7

| |
|---|
| RAD51B |
| ATR |
| WRN |
| RAD18 |
| XRCC5 |
| PRKDC |
| PMS1 |
| TOP3A |
| CCNH |
| DCLRE1B |
| TDP1 |
| POLA1 |
| RAD54L |
| MLH3 |
| ALKBH3 |
| RAD1 |
| XPC |
| POLD1 |
| GTF2H4 |
| NBN |
| RAD51C |
| RAD52 |
| DUT |
| TOP3B |
| CDK7 |
| RFC2 |
| CRY2 |
| EXO1 |
| APTX |
| RAD23A |
| ERCC4 |
| RFC5 |
| XRCC3 |
| ALKBH2 |

Table S1. Core enrichment genes in Hallmark_Oxidative_Phosphorylation for Cdc73-induced t

| RANK | SYMBOL |
|---|---|
| 1 | MTRF1 |
| 2 | NDUFB3 |
| 3 | COX7A2L |
| 4 | NDUFB4 |
| 5 | TOMM70 |
| 6 | ECHS1 |
| 7 | FH |
| 8 | MDH1 |
| 9 | NDUFAB1 |
| 10 | ACADSB |
| 11 | VDAC3 |
| 12 | DECR1 |
| 13 | OPA1 |
| 14 | NDUFB1 |
| 15 | ATP5F1C |
| 16 | OAT |
| 17 | ABCB7 |
| 18 | HSPA9 |
| 19 | AIFM1 |
| 20 | SUPV3L1 |
| 21 | MRPS22 |
| 22 | ATP6V1G1 |
| 23 | NDUFV2 |
| 24 | PRDX3 |
| 25 | ACADM |
| 26 | NDUFA6 |
| 27 | IDH1 |
| 28 | IMMT |
| 29 | ETFA |
| 30 | SLC25A12 |
| 31 | HADHA |
| 32 | NDUFS1 |
| 33 | FXN |
| 34 | SDHC |
| 35 | MDH2 |
| 36 | NDUFB2 |
| 37 | ATP5PB |
| 38 | NDUFS4 |
| 39 | ATP5PF |
| 40 | UQCRC2 |
| 41 | ACAA2 |
| 42 | COX10 |
| 43 | MRPL15 |

| | |
|---|---|
| 44 | SDHB |
| 45 | RETSAT |
| 46 | TIMM9 |
| 47 | MRPL11 |
| 48 | DLD |
| 49 | UQCRB |
| 50 | ETFDH |
| 51 | CYB5R3 |
| 52 | ALDH6A1 |
| 53 | ATP6V1D |
| 54 | HADHB |
| 55 | NDUFA8 |
| 56 | CYB5A |
| 57 | IDH3B |
| 58 | FDX1 |
| 59 | UQCRFS1 |
| 60 | NDUFS3 |
| 61 | NDUFA2 |
| 62 | SUCLG1 |
| 63 | NDUFS2 |
| 64 | NQO2 |
| 65 | NDUFV1 |
| 66 | CPT1A |
| 67 | ATP5PD |
| 68 | RHOT1 |
| 69 | AFG3L2 |
| 70 | NDUFS7 |
| 71 | ACAT1 |
| 72 | PDP1 |
| 73 | UQCR10 |
| 74 | MTX2 |
| 75 | ATP5MF |
| 76 | ATP6V1C1 |
| 77 | COX5A |
| 78 | NDUFS6 |
| 79 | MRPL35 |
| 80 | UQCRQ |
| 81 | PDHX |
| 82 | MTRR |
| 83 | TIMM17A |
| 84 | VDAC1 |
| 85 | COX6B1 |
| 86 | COX7C |
| 87 | NDUFA9 |
| 88 | OXA1L |

Table S1. Core enrichment genes in Hallmark_Oxidative_Phosphorylation for Cdc73-induced t

| RANK | SYMBOL |
|---|---|
| 1 | IDH1 |
| 2 | ECHS1 |
| 3 | NDUFB4 |
| 4 | PRDX3 |
| 5 | MTRF1 |
| 6 | MDH1 |
| 7 | ACADM |
| 8 | NDUFA6 |
| 9 | ACADSB |
| 10 | SDHC |
| 11 | IDH3B |
| 12 | FH |
| 13 | NDUFAB1 |
| 14 | NDUFS7 |
| 15 | CYB5R3 |
| 16 | CPT1A |
| 17 | TOMM70 |
| 18 | OAT |
| 19 | NDUFB3 |
| 20 | MDH2 |
| 21 | SUPV3L1 |
| 22 | ATP5F1C |
| 23 | SDHB |
| 24 | NDUFS2 |
| 25 | COX7A2L |
| 26 | RETSAT |
| 27 | NDUFV1 |
| 28 | HADHA |
| 29 | ACAA2 |
| 30 | AIFM1 |
| 31 | IMMT |
| 32 | ATP6V1G1 |
| 33 | CYB5A |
| 34 | NDUFV2 |
| 35 | FXN |
| 36 | NDUFB1 |
| 37 | NDUFB2 |
| 38 | NDUFA3 |
| 39 | OPA1 |
| 40 | ALDH6A1 |
| 41 | SUCLG1 |
| 42 | ETFDH |
| 43 | MRPL11 |

| Table S2: Flow cytomery reagent | Color | Manufacturer | Cat Number | Clone |
|---|---|---|---|---|
| **Mouse** | | | | |
| B220 | FITC | Biolegend | 103205 | RA3-6B2 |
| 7-AAD | 7-AAD | Biolegend | 420403 | N/A |
| annexin-V | APC | Biolegend | 640920 | N/A |
| c-Kit | APC-Cy7 | Biolegend | 105825 | 2B8 |
| c-Kit | PE | Biolegend | 105807 | 2B8 |
| CD11b | APC | Biolegend | 101211 | M1/70 |
| CD11b | FITC | Biolegend | 101205 | M1/70 |
| CD11b | APCcy7 | Biolegend | 101225 | M1/70 |
| CD11c | FITC | Biolegend | 117306 | N418 |
| CD19 | FITC | BD Biosciences | 553785 | ID3 |
| CD19 | PEcy7 | Biolegend | 115519 | 6D5 |
| CD25 | PE | Biolegend | 102007 | PC61 |
| CD25 | BV605 | Biolegend | 102035 | PC61 |
| CD27 | PE-Cy7 | Biolegend | 124215 | LG.3A10 |
| CD28 | FITC | Biolegend | 122010 | E18 |
| CD3 | FITC | Biolegend | 100305 | 145-2C11 |
| CD4 | FITC | Biolegend | 100539 | RM4-5 |
| CD4 | FITC | Biolegend | 100515 | RM4-5 |
| CD44 | FITC | Biologend | 103011 | IM7 |
| CD45 | FITC | Biolegend | 304011 | H130 |
| CD8a | FITC | Biolegend | 100705 | 53-6.7 |
| CD8b | FITC | Biolegend | 140403 | 53-5.8 |
| CellRox Deep Red | FITC | Invitrogen | C10422 | N/A |
| Flt3 | FITC | Biolegend | 135309 | A2F10 |
| Gr1 | FITC | Biolegend | 108407 | RB6-8C5 |
| Gr1 | FITC | Biolegend | 108405 | RB6-8C5 |
| Gr1 | FITC | Biolegend | 108415 | RB6-8C5 |
| IgM | FITC | eBioscience | 12-5790-82 | II/41 |
| IL7Ra | FITC | Biolegend | 135013 | A7R34 |
| Ly6d | FITC | Biolegend | 138605 | 49-H4 |
| CD45 | FITC | Biolegend | 103131 | 30-F11 |
| NK1.1 | FITC | Biolegend | 108706 | PK136 |
| Sca-1 | FITC | Biolegend | 122523 | E13-161.7 |
| TCRb | FITC | Biolegend | 109206 | H57-597 |
| TCRd | FITC | eBioscience | 11-5711-82 | eBioGL3 |
| Ter119 | FITC | eBioscience | 11-5921-82 | TER-119 |
| Tetramethylrhodamine, Methyl Ester, Perchlorate (TMRM) | FITC | Invitrogen | T668 | N/A |
| **Human** | | | | |
| CD7 | APC | Biolegend | 343107 | 4H9 |
| CD45 | APC | Biolegend | 304011 | H130 |
| LNGFR (CD271) | APC | Miltenyl Biotech | 130-091-884 | ME20.4-1.4H |

| Table S2: Antibody | Use | Manufacturer | Cat Number | Clone |
| --- | --- | --- | --- | --- |
| Ets1 | Western Blot/Co-IP | Cell Signaling Technologies | 14069S | D8O8A |
| Rabbit IgG | Co-Immunoprecipitation | Cell Signaling Technologies | 2729S | N/A |
| ECL Rabbit IgG, HRP-linked whole Ab (from donkey) | Westen Blot Secondary Ab | GE Healthcare Life Sciences | NA934 | N/A |
| ECL Mouse IgG, HRP-linked whole Ab (from sheep) | Westen Blot Secondary Ab | GE Healthcare Life Sciences | NA931 | N/A |
| Peroxidase-conjugated AffiniPure Rabbit Anti-Sheep IgG | Western Blot Secondary Ab | Jackson ImmunoResearch | 313-035-003 | N/A |
| CDC73 | Western Blot | R&D Systems | AF5508 | N/A |
| anti-Parafibromin (Cdc73) | ChIP-Seq/Co-IP/Western Blot | Bethyl | A300-171A (discontinued) | N/A |
| anti-Rabbit IgG, peroxidase-linked species-specific whole antibody (from donkey) | Western Blot Secondary Ab | GE Healthcare Life Sciences | NA934 | N/A |
| b-actin | Western Blot | Sigma | A5316 | N/A |
| Ets1 | ChIP-Seq | Santa Cruz | sc-350x (discontinued) | N/A |
| GAPDH | Western Blot | Cell Signaling Technologies | 5174 | D16H11 |
| Goat anti-Mouse IgG (H+L), HRP | Western Blot Secondary Ab | Invitrogen | 31430 | N/A |
| H3K27Ac | ChIP-Seq | Active Motif | 39133 | N/A |
| Histone H2A.XS139ph | ChIP | Active Motif | 39117/8 | N/A |
| Phospho-Chk1 (Ser296) | Western Blot | Cell Signaling Technologies | 90178 | D3O9F |
| Phospho-Histone H2A.X (Ser139) | Western Blot | Cell Signaling Technologies | 9718 | 20E3 |
| Rabbit Anti-Goat IgG, HRP conjugate | Western Blot Secondary Ab | Millipore Sigma | AP106P | N/A |
| Ubiquityl-Histone H2B (Lys120) | ChIP-Seq | Cell Signaling Technologies | 5546 | D11 |

**Table S2: Primers**

| Mouse qRT-PCR primers | Forward (5' to 3') | Reverse (5' to 3') |
|---|---|---|
| m18S | GCGCCGCTAGAGGTGAAAT | GGCGGGTCATGGGAATAAC |
| mEf1a | CACTTGGTCGCTTTGCTGTT | GGTGGCAGGTGTTAGGGGTA |
| mEts1 | CAAGCCGACTCTCACCATCA | ATTCCCAGTCGCTGCTGTTC |
| mAtr | GCTGGCCACCATCAGACAGC | ACGTCACCCTTGGACCACAGC |
| mB-actin | GCCCTGAGGCTCTTTTCCAG | TGCCACGGATTCCATACC |
| mCdc73 | CCGGAAAGGAAGGCCAACCCA | AGCTGCACGCCGGACATAAACA |
| mHes1 | GGAAATGACAGTGAAGCACCT | CAGCACACTTGGGTCTGTG |
| mLig4 | GGGTGACTTGGAGCAGCTGAGG | TGGAGATGGGCTTCCGCCTT |
| mMyc | Taqman Mm00487803_m1 | |
| mNdufab1 | GTGAAACCCACACTGCTGTT | AGGTACCGTCTCTCTGGTCT |
| mNdufb4 | ACCCTCGACCCTGCCGAGTA | CCCGTTTAAGCCGGGCCCTT |
| **Mouse ChIP primers** | **Forward (5' to 3')** | **Reverse (5' to 3')** |
| Fgg | GGTGGCAGATGGGAGGGAAC | CCAGGGGTGGGAGAAGGGAA |
| mCol1a1 | TGCAGACAAGCCCCTCAGTG | ACGAAGGTGGCATGAAGGAAC |
| mHbaa1 | TTCCTCATGCAAAGCGAACA | CCTGGTAGGCTGAGGCAAGA |
| mLig4 | GGGTGACTTGGAGCAGCTGAG | TGGAGATGGGCTTCCGCCTT |
| mMyc | GCATAGACCTCATCTGCGGTTG | AAGGGGGAAGGACGAACGAATG |
| **Genotyping primers** | **Forward (5' to 3')** | **Reverse (5' to 3')** |
| LckCre | CCTCCTGTGAACTTGGTGCTTGAG | TGCATCGACCGGTAATGCAG |
| Cdc73 | TCCTTTCCATTGTGCAGCTGGTTG | TGCCAGTGCAAGAACCTCATCCTA |
| Rosa26-CreERT2 | AAAGTCGCTCTGAGTTGTTAT | CCTGATCCTGGCAATTTCG |
